# LY-CoV1404 (bebtelovimab) potently neutralizes SARS-CoV-2 variants

**DOI:** 10.1101/2021.04.30.442182

**Authors:** Kathryn Westendorf, Stefanie Žentelis, Lingshu Wang, Denisa Foster, Peter Vaillancourt, Matthew Wiggin, Erica Lovett, Robin van der Lee, Jörg Hendle, Anna Pustilnik, J. Michael Sauder, Lucas Kraft, Yuri Hwang, Robert W. Siegel, Jinbiao Chen, Beverly A. Heinz, Richard E. Higgs, Nicole L. Kallewaard, Kevin Jepson, Rodrigo Goya, Maia A. Smith, David W. Collins, Davide Pellacani, Ping Xiang, Valentine de Puyraimond, Marketa Ricicova, Lindsay Devorkin, Caitlin Pritchard, Aoise O’Neill, Kush Dalal, Pankaj Panwar, Harveer Dhupar, Fabian A. Garces, Courtney A. Cohen, John M. Dye, Kathleen E. Huie, Catherine V. Badger, Darwyn Kobasa, Jonathan Audet, Joshua J. Freitas, Saleema Hassanali, Ina Hughes, Luis Munoz, Holly C. Palma, Bharathi Ramamurthy, Robert W. Cross, Thomas W. Geisbert, Vineet Menacherry, Kumari Lokugamage, Viktoriya Borisevich, Iliana Lanz, Lisa Anderson, Payal Sipahimalani, Kizzmekia S. Corbett, Eun Sung Yang, Yi Zhang, Wei Shi, Tongqing Zhou, Misook Choe, John Misasi, Peter D. Kwong, Nancy J. Sullivan, Barney S. Graham, Tara L. Fernandez, Carl L. Hansen, Ester Falconer, John R. Mascola, Bryan E. Jones, Bryan C. Barnhart

## Abstract

SARS-CoV-2 neutralizing monoclonal antibodies (mAbs) can reduce the risk of hospitalization when administered early during COVID-19 disease. However, the emergence of variants of concern has negatively impacted the therapeutic use of some authorized mAbs. Using a high throughput B-cell screening pipeline, we isolated a highly potent SARS-CoV-2 spike glycoprotein receptor binding domain (RBD)-specific antibody called LY-CoV1404 (also known as bebtelovimab). LY-CoV1404 potently neutralizes authentic SARS-CoV-2 virus, including the prototype, B.1.1.7, B.1.351 and B.1.617.2). In pseudovirus neutralization studies, LY-CoV1404 retains potent neutralizing activity against numerous variants including B.1.1.7, B.1.351, B.1.617.2, B.1.427/B.1.429, P.1, B.1.526, B.1.1.529, and the BA.2 subvariant and retains binding to spike proteins with a variety of underlying RBD mutations including K417N, L452R, E484K, and N501Y. Structural analysis reveals that the contact residues of the LY-CoV1404 epitope are highly conserved with the exception of N439 and N501. Notably, the binding and neutralizing activity of LY-CoV1404 is unaffected by the most common mutations at these positions (N439K and N501Y). The breadth of reactivity to amino acid substitutions present among current VOC together with broad and potent neutralizing activity and the relatively conserved epitope suggest that LY-CoV1404 has the potential to be an effective therapeutic agent to treat all known variants causing COVID-19.

**In Brief:** LY-CoV1404 is a potent SARS-CoV-2-binding antibody that neutralizes all known variants of concern and whose epitope is rarely mutated.

**Highlights:**

- LY-CoV1404 potently neutralizes SARS-CoV-2 authentic virus and known variants of concern including the B.1.1.529 (Omicron), the BA.2 Omicron subvariant, and B.1.617.2 (Delta) variants
- No loss of potency against currently circulating variants
- Binding epitope on RBD of SARS-CoV-2 is rarely mutated in GISAID database
- Breadth of neutralizing activity and potency supports clinical development

## Introduction

Variants of SARS-CoV-2 continue to alter the trajectory of the COVID-19 pandemic, which at the time of writing has infected over 416 million people world-wide and is responsible for more than 5.8 million deaths (https://covid19.who.int/ accessed 17 February 2022). As predicted, SARS-CoV-2 has continued to evolve as the pandemic has progressed (Mercatelli and Giorgi, 2020; Pachetti et al., 2020). Selective pressures and viral adaptation during prolonged, suboptimally-treated infections are thought to have generated numerous variants (McCormick et al., 2021), some significantly diminishing the effectiveness of COVID-19 clinical countermeasures (Altmann et al., 2021; Plante et al., 2020). Variants of concern (VOC) represent a closely monitored subset of the many detected SARS-CoV-2 variants, due to their potential for increased transmissibility, and ability to evade immunity produced by infection or vaccines while reducing the efficacies of antibody-based treatments (Hoffmann et al., 2021; Kuzmina et al., 2021; McCormick et al., 2021; Thomson et al., 2021; Wang et al., 2021b). The impact of VOC continues to increase (World Health Organization, 2021), with emerging variants threatening to slow the pace and success of global vaccination efforts and limit the effectiveness of existing COVID-19 treatments (Davies et al.; Munnink et al., 2021; Plante et al., 2020; Tegally et al., 2021).

Many studies have demonstrated the clinical safety and efficacy of antibody-based COVID-19 therapies and their potential for easing the strain on economies and healthcare infrastructures during the pandemic (Chen et al., 2020; Jiang et al., 2020; Jones et al., 2020). Key target populations for such monoclonal antibody treatments include immunocompromised individuals, patients that are particularly susceptible due to comorbidities (cardiovascular disease, diabetes, chronic respiratory conditions), and those aged over 65 (Jiang et al., 2020). To date, several antibody therapies targeting the spike (S) protein of SARS-CoV-2 have been successfully used in the fight against the virus (Chen et al., 2020; Gottlieb et al., 2021). One example is Eli Lilly and Companies’ bamlanivimab (LY-CoV555), which was the first anti-SARS-CoV-2 monoclonal antibody tested clinically in June 2020, three months after initiating discovery efforts (Jones et al., 2021). Lilly’s combination of bamlanivimab with another antibody, etesevimab, received emergency authorization in several countries. Clinical testing has shown that antibodies from Regeneron, GSK/Vir, AstraZeneca, and others are also safe and effective (AstraZeneca, 2020; GlaxoSmithKline, 2021; Regeneron, 2021; Tada, et al., 2021). As variants continue to emerge and spread, antibodies can provide highly effective therapeutic options to vulnerable populations against COVID-19. However, for antibody therapies to be successful, they must retain potent neutralizing breadth against emerging SARS-CoV-2 lineages carrying mutations. A high neutralization potency may also provide the opportunity to explore lower clinical doses (Andreano et al., 2021) and alternate routes of administration. With the rise of Omicron and BA.2 VOC many of the authorized antibodies may no longer be an option with a strong negative impact on their neutralization capacity (Iketani et al., 2022).

Multiple reports have consistently shown that mutations in SARS-CoV-2 can substantially reduce the binding affinity and neutralization of antibodies (Rees-Spear et al., 2021; Yao et al., 2021). Variants such as B.1.1.529, BA.2, B.1.1.7, B.1.351, P.1, B.1.526, B.1.427, and B.1.429 affect the *in vitro* binding of antibodies being tested clinically or on those already authorized for emergency use to varying degrees (Kuzmina et al., 2021; Liu et al., 2021; Rees-Spear et al., 2021; Iketani et al., 2022). Mutations at amino acid positions 417, 439, 452, 484, and 501 were found to have the greatest consequences on the functionality of both antibodies and vaccines (Thomson et al., 2021). Therefore, antibodies that retain binding and potent neutralization activity in the presence of these mutations could be extremely valuable tools to rapidly respond to these variants.

In this report, we describe LY-CoV1404 (also known as bebtelovimab), a fully human IgG1 monoclonal SARS-CoV-2 antibody targeting the receptor binding domain (RBD), identified in our ongoing pandemic-response efforts. As of February 2022, LY-CoV1404 binds and potently neutralizes all currently known VOC of SARS-CoV-2 including B.1.1.529 and BA.2 (the Omicron variant and sub-variant). LY-CoV1404 binds to an epitope that is largely distinct from the mutations identified to be widely circulating within the newly emerged variants, including those mutations that reduce the effectiveness of vaccines. Importantly, the LY-CoV1404 interaction with the S protein is driven by amino acids that are rarely mutated in the global GISAID EpiCoV database, indicating LY-CoV1404 has the potential to be a long-term, solution for reducing COVID-19-related illness and death. Additionally, bebtelovimab received Emergency Use Authorization from the US FDA on February 11, 2022 (https://www.fda.gov/news-events/press-announcements/coronavirus-covid-19-update-fda-authorizes-new-monoclonal-antibody-treatment-covid-19-retains).

## Results

### Monoclonal antibody LY-CoV1404 retains binding to known variants of SARS-CoV-2

Variants of SARS-CoV-2 continue to be identified globally (**Figure 1A**) and can abrogate the binding of monoclonal antibody therapies or reduce vaccine effectiveness (Hoffmann et al., 2021; Wang et al., 2021b). To identify potent neutralizing antibodies to SARS-CoV-2 variants, a high-throughput screening approach (Jones et al., 2021) was conducted on peripheral blood mononuclear cells (PBMC) that were isolated from a COVID-19 convalescent donor. The sample was obtained approximately 60 days after the onset of symptoms.

**Figure 1.**
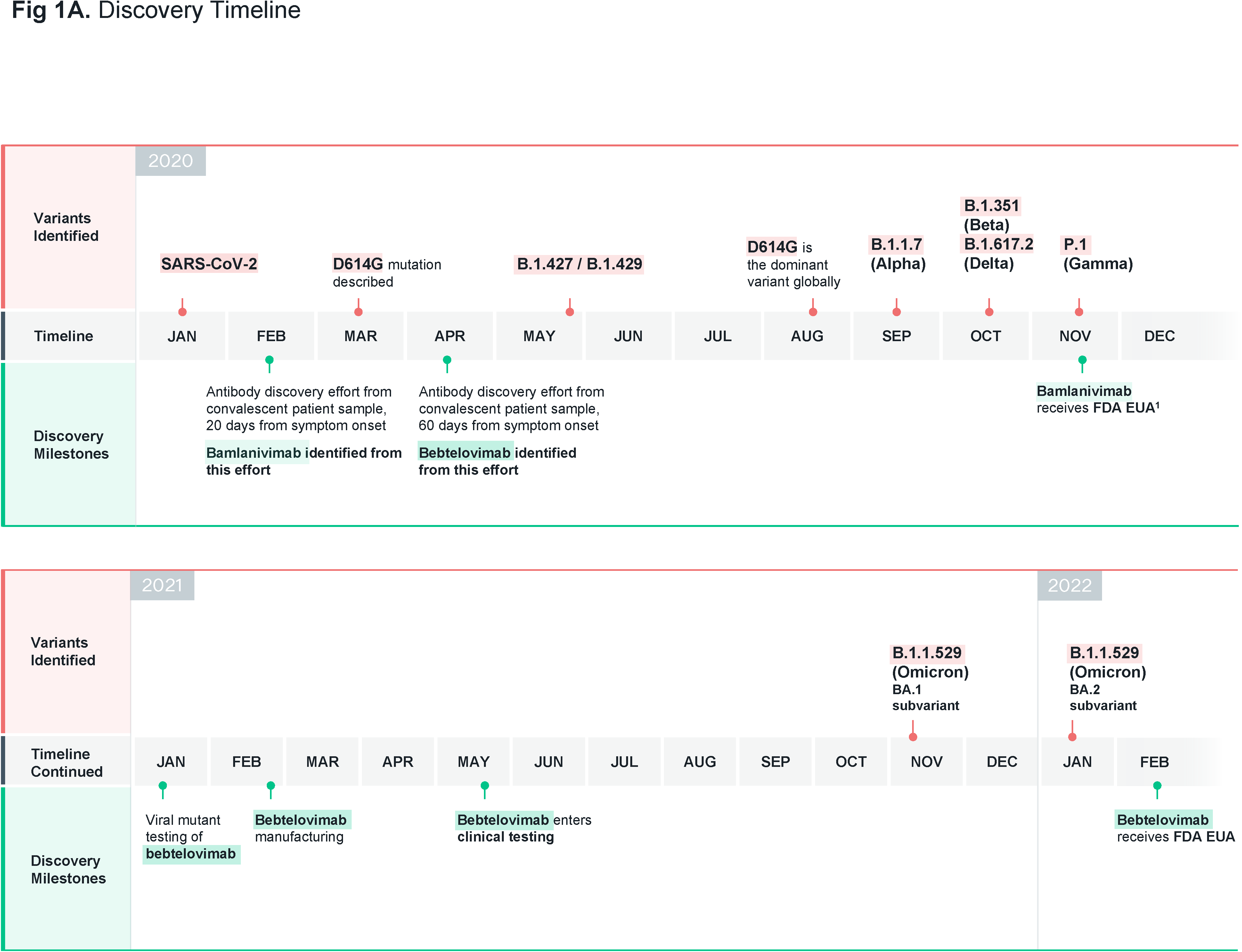
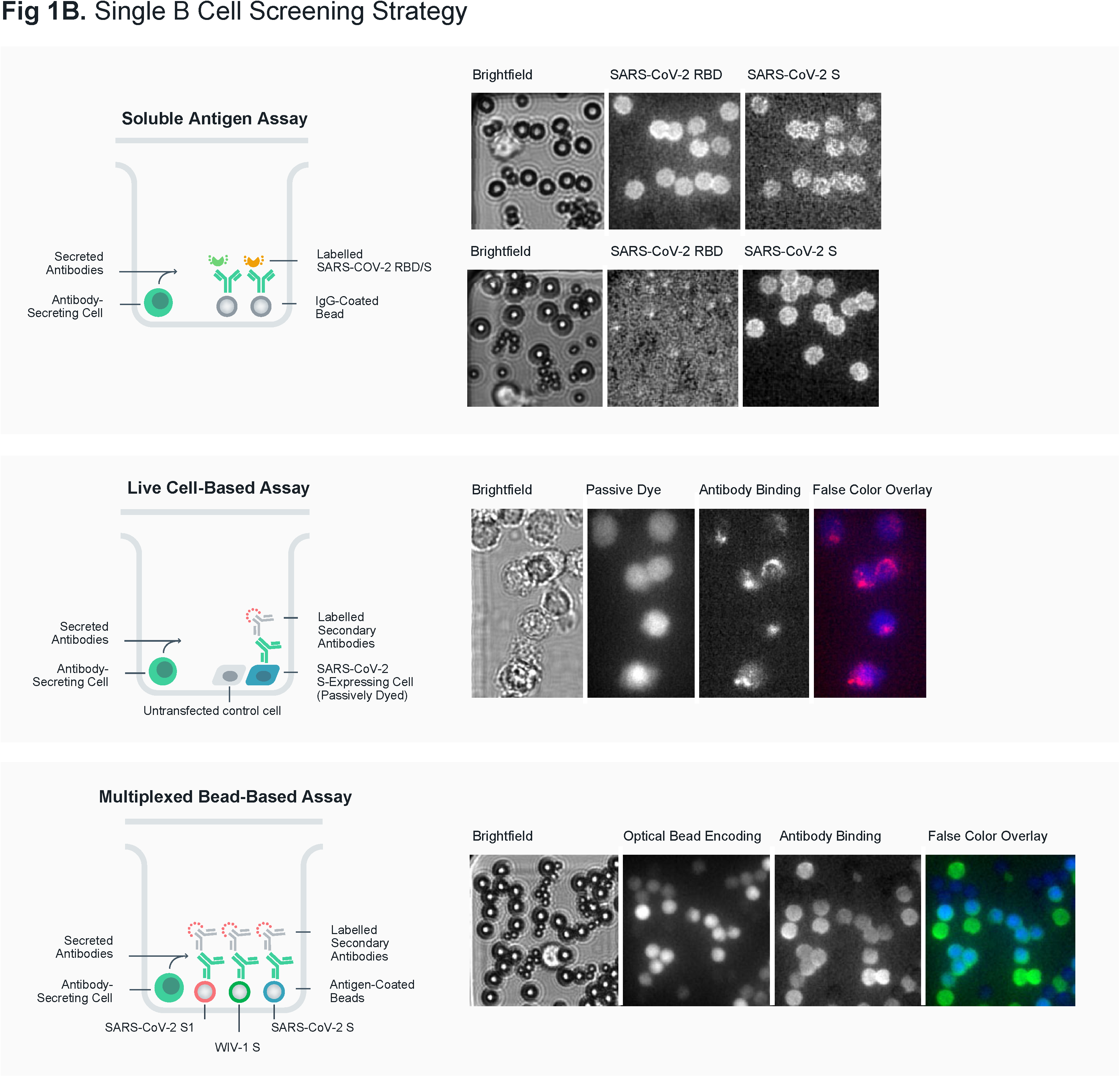
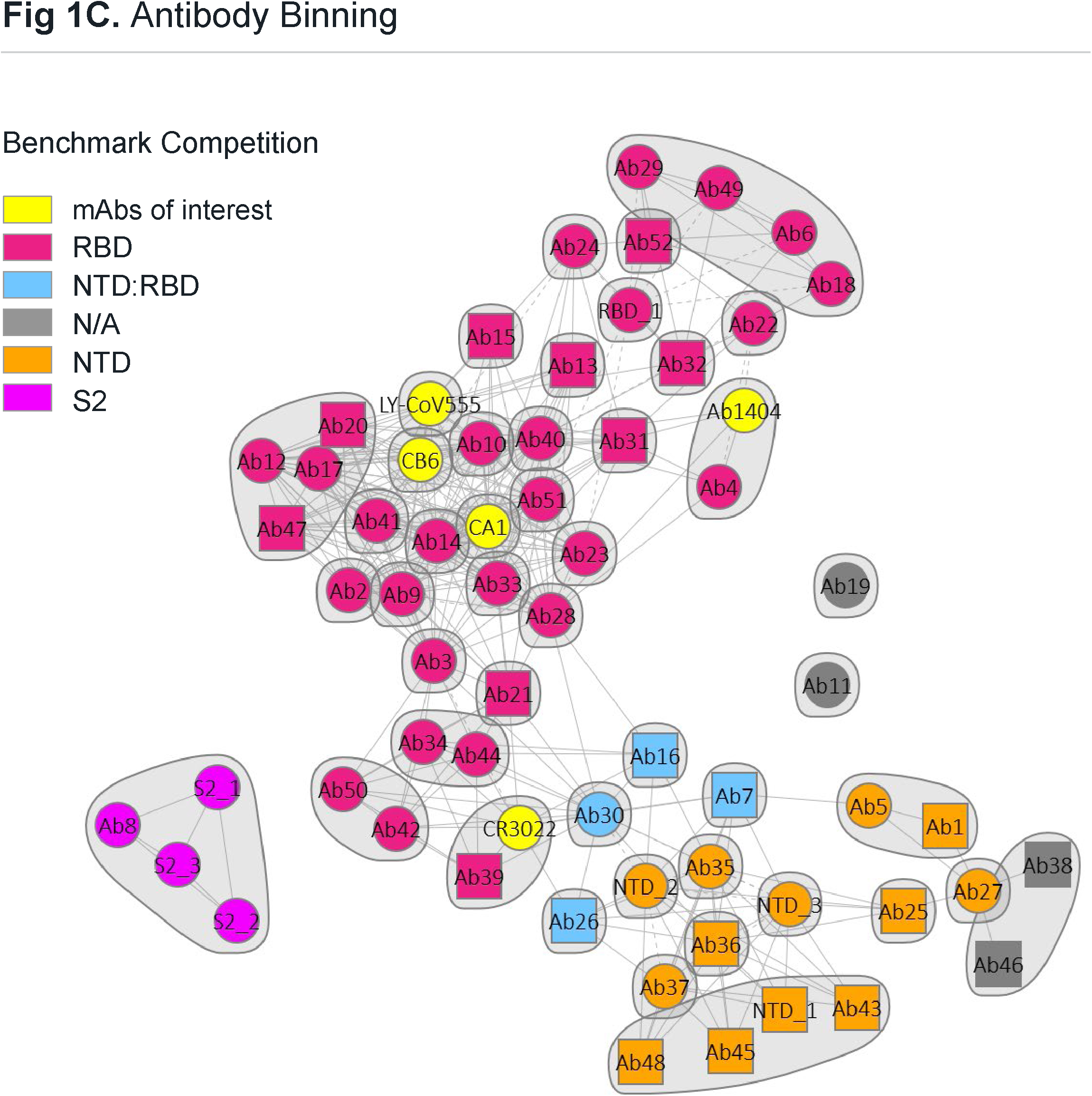
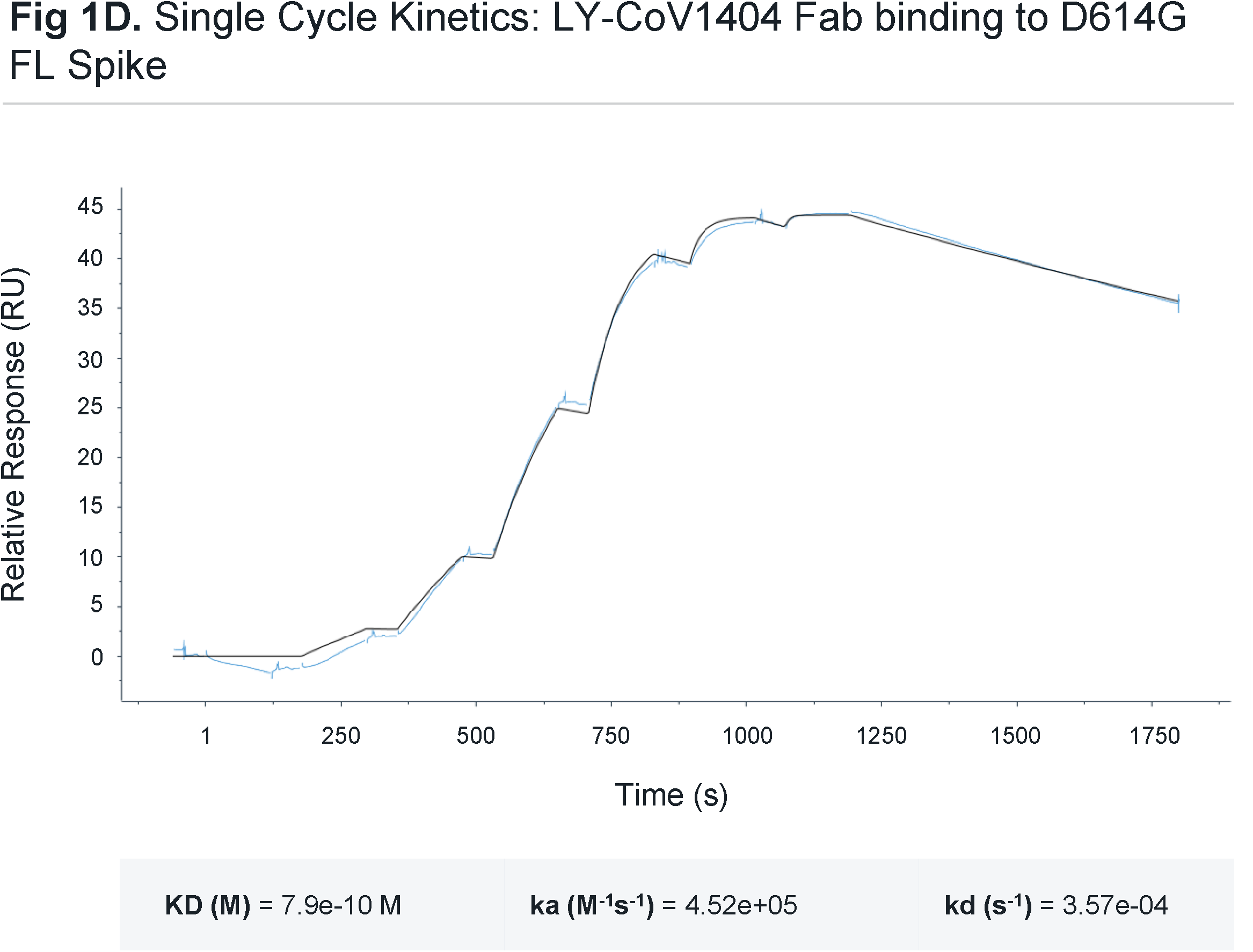
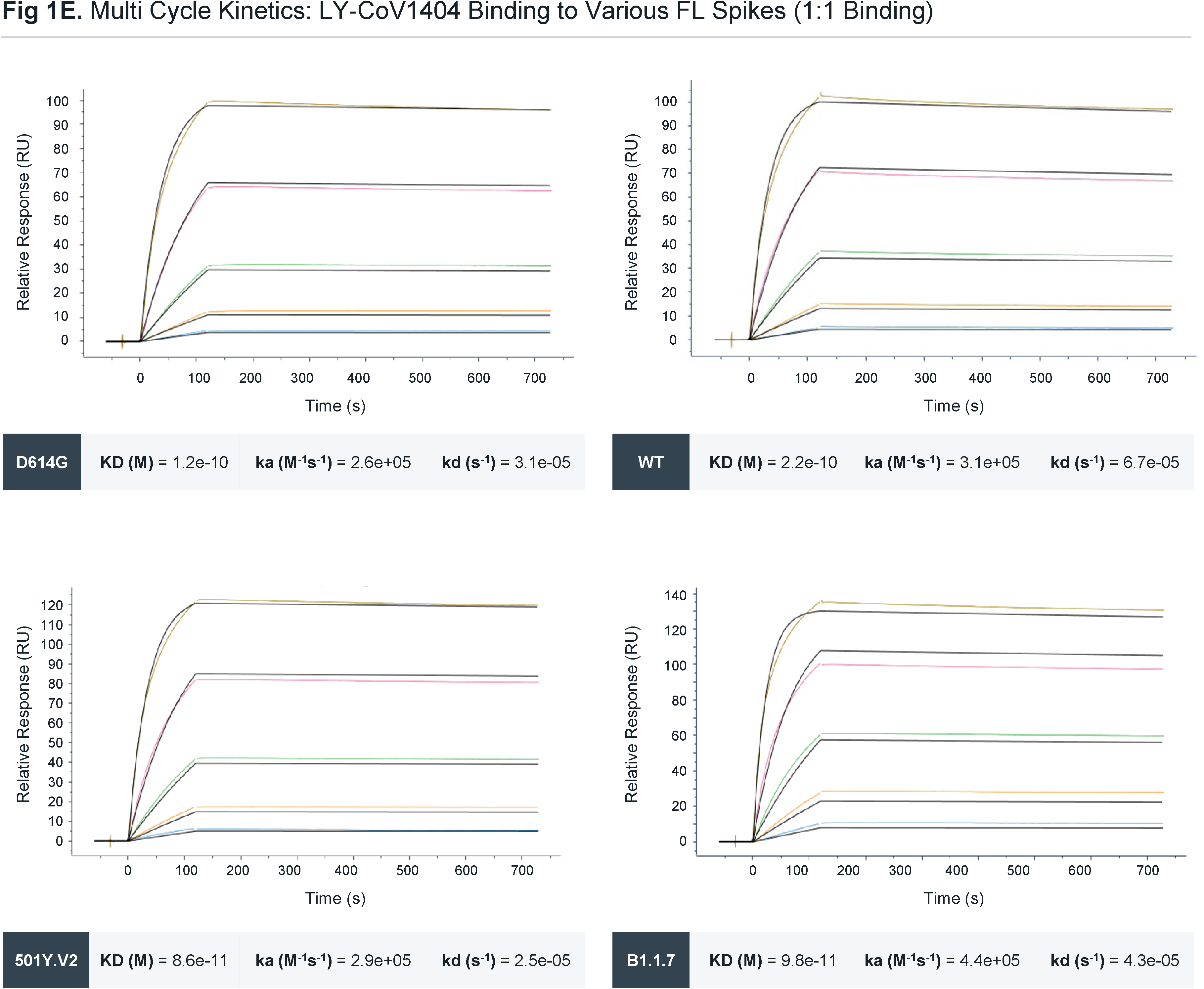
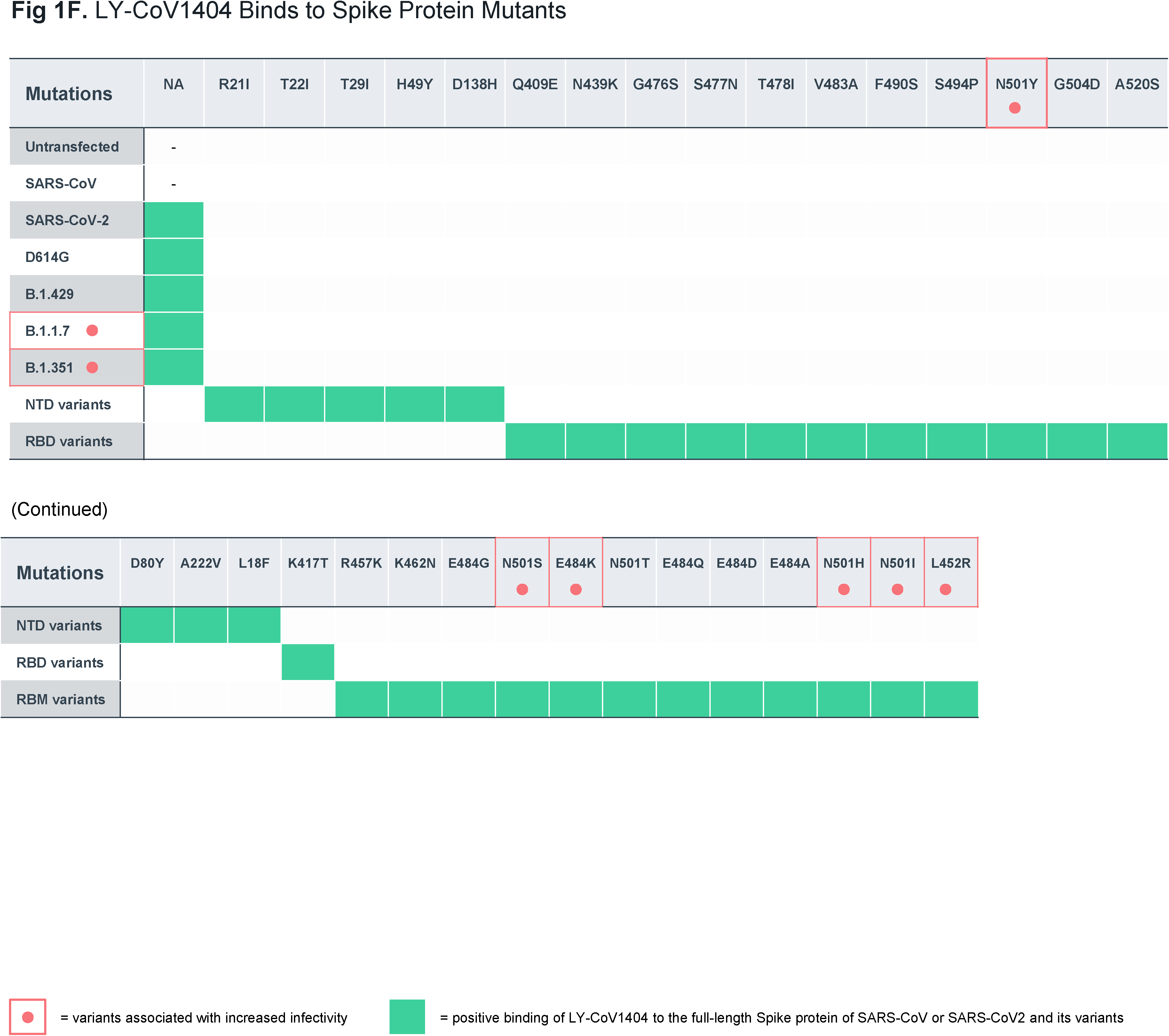
LY-CoV1404 discovery and binding properties. (A) Timeline (top) of identification of Variants of Concern (VOC) to date in the COVID-19 pandemic. Milestones of discovery of antibody therapies to treat SARS-CoV-2 infection (bottom). (B) Representation of soluble antigen assay, live cell-based screening assay and multiplexed bead-based assay. Representative microscopic images of antibodies assessed for SARS-CoV-2 spike protein specificity in each indicated assay. (C) Epitope binning and isolated subdomain binding for discovered antibodies and benchmarks. Each antibody was tested in two orientations: as a ligand on the chip, and as an analyte in solution. Individual antibodies are represented either as a circle (data present in both orientations) or as a square (data present with the antibody in a single orientation). Bins are represented as envelopes (46 total) and competition between antibodies as solid (symmetric competition) or dashed (asymmetric competition) lines. Benchmark-based blocking profiles are indicated by color. (D) Single cycle kinetics of LY-CoV1404 Fab run on full-length spike protein carrying the D614G mutation. (E) Multi-cycle kinetics of LY-CoV1404 run on indicated full length spike protein. (F) Binding of LY-CoV1404 to indicated spike proteins expressed on CHO cells.

Three different screening strategies were employed to identify SARS-CoV-2-binding antibodies from the PBMCs in this donor sample. We used a soluble antigen assay with fluorescently labeled prefusion-stabilized full-length trimeric SARS-CoV-2 S protein as well as the SARS-CoV-2 RBD subunit, a live-cell-based assay using mammalian cells that transiently expressed the SARS-CoV-2 S protein, and a multiplexed bead assay using S proteins from multiple coronaviruses (**Figure 1B**). A total of 740,000 cells were screened in this discovery effort and machine learning (ML)-based analyses were used to automatically select and rank 1,692 antibody “hits” (0.22% frequency). Of these, 1,062 single antibody-secreting cells were selected for recovery. Libraries of antibody genes from recovered single B cells were generated and sequenced using next-generation sequencing, and a custom bioinformatics pipeline with ML-based sequence curation was used to identify paired-chain antibody sequences, with 290 unique high-confidence paired heavy and light chain sequences identified (**Figure 1B**). The antibody sequences corresponded to 263 clonal families and used a diverse set of 41 VH genes. The mean sequence identity to germline was 96.8% from this 60-days post-symptom-onset blood draw in this donor.

High-throughput SPR experiments on a selected subset of 69 recombinantly-expressed antibodies (including benchmark controls) were performed to assess S protein epitope coverage. These studies included epitope binning, isolated subdomain binding, and binding competition with ACE2 (**Figures 1C**/**S1**). We employed 21 benchmark antibodies including those known to bind to S protein subunits S1, N-terminal domain (NTD), RBD, S2 subunit, and several internally identified antibodies with known binding domains. The antibodies (including benchmarks) were clustered into 46 bins using Carterra epitope analysis software based on the competition heatmap (**Figure 1C**) and were delineated into bins among known subunits of the S protein. Several of the antibodies discovered are cross-blocked with those that are currently in clinical testing or authorized for use in COVID-19 patients including bamlanivimab. We therefore focused analyses on antibodies outside of these established epitopes and identified LY-CoV1404, an ACE2-blocking antibody, which is found in a unique bin shared with one other antibody. LY-CoV1404 was observed to compete directly for S protein RBD binding with the S309 (Pinto et al., 2020) and REGN10987 (Hansen et al., 2020) antibodies, suggesting a binding epitope somewhat distinct from other ACE2-blocking antibodies such as bamlanivimab or etesevimab.

Characterization of LY-CoV1404 binding kinetics revealed that the Fab fragment of the antibody bound to the S protein of D614G with high affinity, characterized by binding constant (K_D_) values ranging between 790 pM and 4nM, depending on the assay (**Figure 1D**, **Table 1**). We further analyzed the binding affinity of LY-CoV1404 as a human IgG1 antibody and determined the K_D_ to be between 75 pM and 220 pM (depending on the assay design) for LY-CoV1404 binding to the S protein (**Figure 1E**, **Table 1**). There was no loss of binding to the S protein carrying the D614G mutation (Asp^614^→Gly).

**Table 1.**
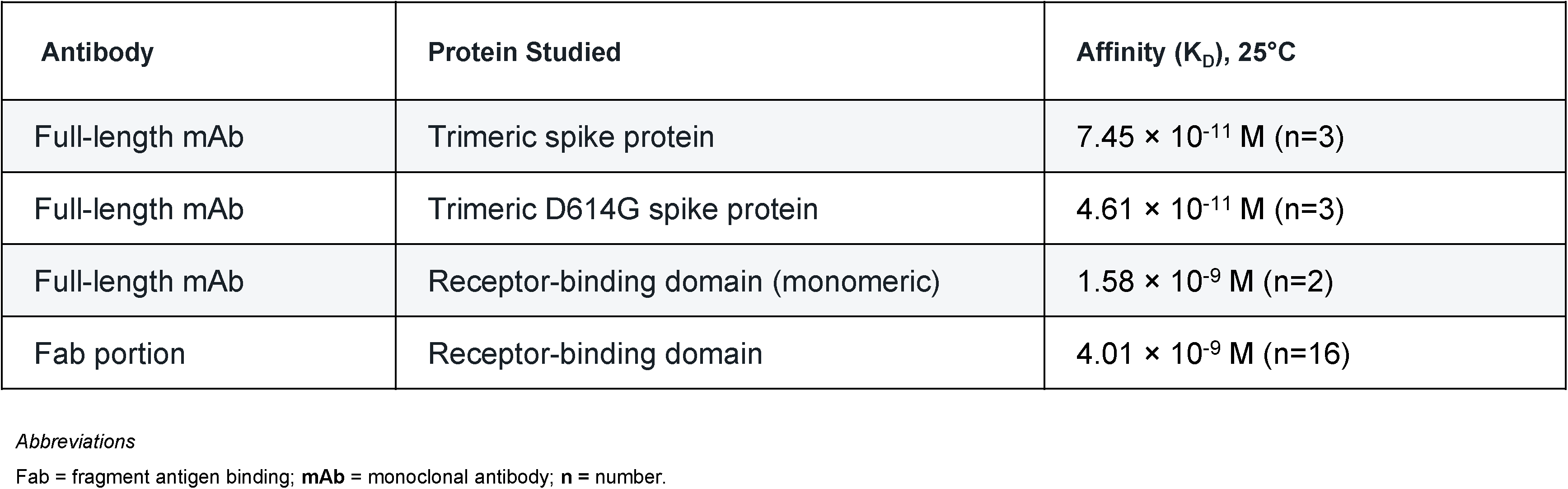
Summary of Target-Binding Affinities Measured through Surface Plasmon Resonance

We explored the ability of LY-CoV1404 to bind variant S proteins from the more widely circulating VOC lineages of SARS-CoV-2. LY-CoV1404 retained binding to key VOC including B.1.351 and B.1.1.7. Critically, when compared to wild type virus, LY-CoV1404 bound to these variants with no loss in affinity (**Figure 1E**). These data suggest that LY-CoV1404 binds to a region that is not significantly affected by the prevalent VOC mutations. We further tested the binding of LY-CoV1404 to a wider range of S mutants and VOC using an assay system with S protein expressed on the surface of mammalian cells. In these assays, we compared binding of LY-CoV1404 to isotype control antibodies. Consistent with the affinity data, LY-CoV1404 maintained binding to all variants and mutants tested (**Figure 1F**). These data indicate that SARS-CoV-2 S protein variants identified and reported to be of significant concern in this pandemic are still potently bound by LY-CoV1404.

### LY-CoV1404 potently neutralizes SARS-CoV-2 authentic virus and viral variants

The *in vitro* viral neutralization activity LY-CoV1404 was assessed against an authentic viral strain (SARS-CoV-2/MT020880.1) using an immunofluorescence assay (IFA) readout. In these studies, LY-CoV1404 potently neutralized the authentic virus with half maximal inhibitory concentration (IC50) values ranging from 9 ng/mL to 22.1 ng/mL. Bamlanivimab (LY-CoV555), another potent anti-RBD binding antibody, consistently had a 2-3-fold higher (less potent) IC50 compared to LY-CoV1404 (**Figure 2A**).

**Figure 2.**
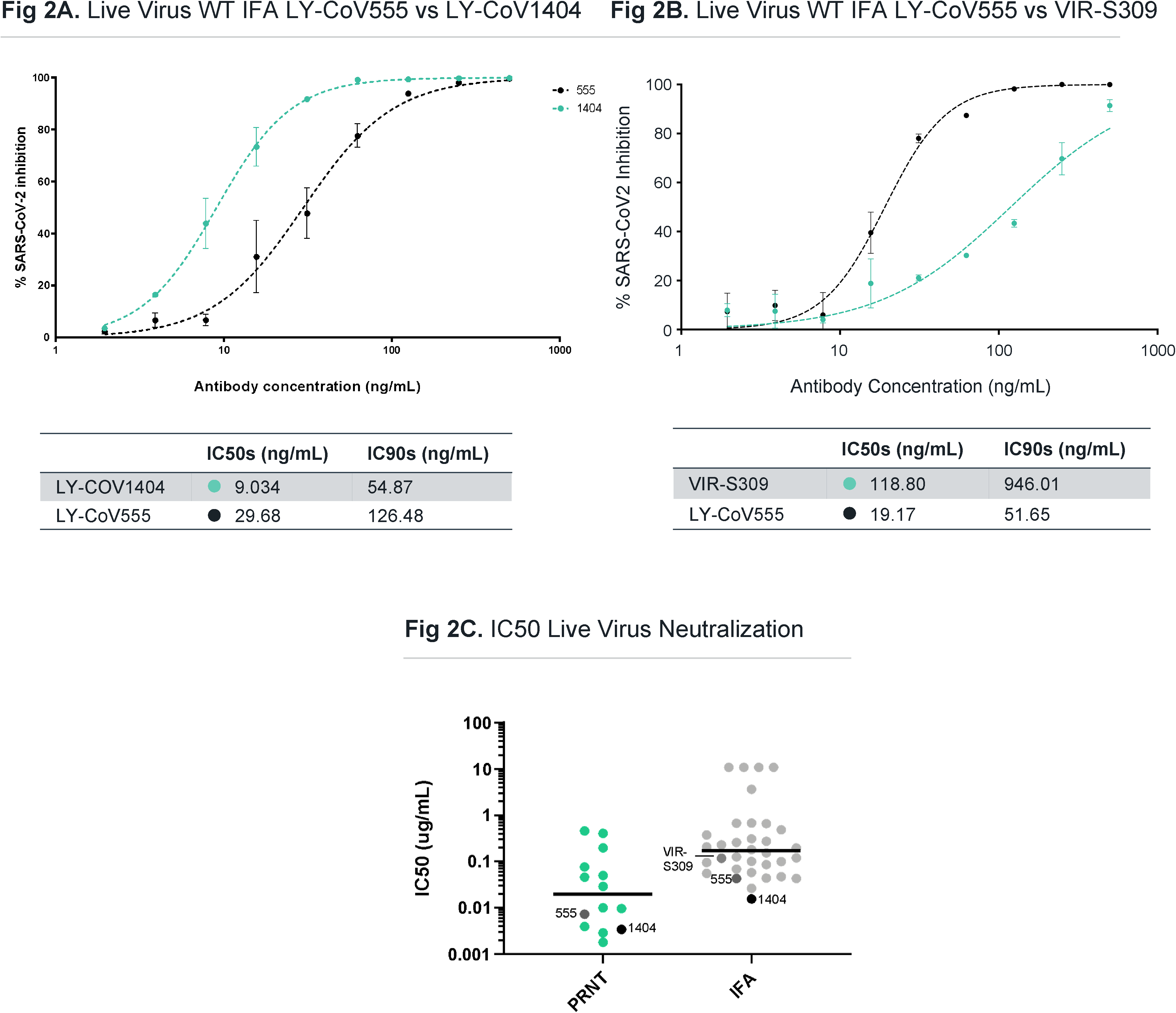
Anti-SARS-CoV-2 monoclonal antibody authentic virus neutralization. (A) Authentic virus (Italy isolate) neutralization using IFA by LY-CoV1404 and LY-CoV555 across a dose range. (B) Authentic virus (Italy isolate) neutralization using IFA by S309 across a dose range. (C) Comparison of authentic virus neutralization by antibodies in PRNT and IFA assays. Specific antibodies are indicated. Data are compiled from several different individual experiments.

We also tested the VIR-S309 antibody that was recently described to have a unique binding epitope, outside of many of the mutated positions in current VOC (Pinto et al., 2020). Vir’s authorized antibody, Vir-7831, is cross-reactive to SARS-CoV-1 S protein (Cathcart et al., 2021). Comparing separate assay runs, we observed VIR-S309 had an IC50 of 118 ng/mL (highly consistent with a recent report on the clinically active derivative VIR-7831 (Cathcart et al., 2021)) compared to bamlanivimab with a range of 19.2 to 29.7 ng/mL IC50 in multiple independent assays (**Figure 2B**). Amongst other antibodies tested, LY-CoV1404 exhibited the most potent neutralization of authentic virus in this assay (**Figure 2C**). We also tested authentic virus neutralization in a plaque reduction neutralization test (PRNT) against a Canadian strain of SARS-CoV-2 (hCoV-19/Canada/ON_ON-VIDO-01-2/2020) with multiple antibodies (**Figure 2C**), and against other natural SARS-CoV-2 isolates tested with LY-CoV1404 alone and in combination with other antibodies (**Table 2**). LY-CoV1404 potently neutralized viral infection in the PRNT assays with IC50 values ranging from 2-10 ng/mL, and there was no impact on neutralization when combined with other neutralizing mAbs.

**Table 2.** Neutralization of Virus Infectivity by Plaque Reduction Assay

|  | IC<sub>50</sub>, µg/mL<br>(95% CI) | IC<sub>90</sub>, µg/mL<br>(95% CI) |
| --- | --- | --- |
| **LY-CoV1404** |  |  |
| WA isolate | 0.002<br>(0.002, 0.004) | 0.0017<br>(0.007, 0.044) |
| Italy-INMI1 | 0.010<br>(0.008, 0.012) | 0.142<br>(0.085, 0.239) |
| Netherlands | 0.005<br>0.004, 0.006) | 0.038<br>(0.022, 0.066) |
| **mAb combinations (molar ratio)\*** |  |  |
| LY-CoV1404 | 0.015<br>(0.012, 0.018) | 0.066<br>(0.038, 0.114) |
| Bamlanivimab | 0.031<br>(0.024, 0.041) | 0.134<br>(0.074, 0.242) |
| Etesevimab | 0.189<br>(0.069, 0.519) | 2.633<br>(0.446, 15.529) |
| LY-CoV1404 + Bamlanivimab (1:2) | 0.030<br>(0.024, 0.037) | 0.128<br>(0.080, 0.206) |
| LY-CoV1404 + Etesevimab (1:4) | 0.049<br>(0.041, 0.058) | 0.185<br>(0.128, 0.267) |
| LY-CoV1404 + Bamlanivimab +<br>Etesevimab (1:2:4) | 0.023<br>(0.018, 0.030) | 0.140<br>(0.076, 0.258) |

*Abbreviations* **CI** = confidence interval. **Absolute IC50** = concentration inhibiting maximal activity by 50%; **IC90** = concentration inhibiting maximal activity by 90%;
\* Combination study was conducted using Italy-INMI1 virus isolate. Data presented as mean of duplicate wells and 95% confidence intervals.

To further characterize the functional activity of LY-CoV1404 against VOCs we tested the neutralization activity in authentic VOC virus assays and pseudotyped neutralization assays with a wider array of VOC and subpopulations. Consistent with the binding data showing high affinity binding to a variety of VOCs and individual mutations, LY-CoV1404 retained full neutralization potency against authentic virus including wild type, B.1.1.7, B.1.351 and B.1.617.2 (**Table 3A**). In pseudotyped neutralization assays using a wide breadth of VOCs, LY-CoV1404 retained activity against all variants, including prominent VOCs B.1.617.2, B.1.351, B.1.1.529, BA.2 and P.1 (**Table 3B**). The rapid spread of the new variant, B.1.1.529 (Omicron), with 35 mutations in the amino acid sequence of the spike protein, and 15 of those mutations located in the RBD, has raised concerns for the activity of a number of clinically tested therapeutic monoclonal antibodies. The ability of these antibodies to neutralize B.1.1.529 was compared in pseudotyped neutralization assays. LY-CoV1404 was the only antibody that retained full potency against the Omicron variant in these assays; separately, Vir-7831 (sotrovimab) has been shown to exhibit a minor loss in neutralization potency (2.7-fold reduction) against this same variant (Cathcart et al., 2021). (**Table 3C**, **3D**). These assays were performed using two different systems with highly comparable results. In addition, we tested the subvariant of B.1.1.529 known as BA.2 in pseudovirus neutralization (**Table 3B**). LY-CoV1404 retained activity against this variant as well. The data indicate that LY-CoV1404 retains its potency against VOCs, including those dominant or rapidly spreading worldwide (such as B.1.617.2, B.1.1.529, and BA.2).

**Table 3A.** Live virus Neutralization of SARS-CoV-2 Variants in the Presence of LY-CoV1404 (PRNT assay)

| SARS-CoV-2 Variant | Viral Isolate | IC50<br>µg/mL<sup>b</sup> | IC90<br>µg/mL<sup>b</sup> |
| --- | --- | --- | --- |
| Wuhan | USA/WA/1/2020<sup>a</sup> | 0.011 | 0.108 |
| Alpha / B.1.1.7 / UK Origin | England/204820464/2020 | 0.004 | 0.044 |
| Beta / B.1.351 / South Africa Origin | USA/MD-HP01542/2021 | 0.007 | 0.045 |
| Delta/B.1.617.2 | GNL-751-TVP23523 | 0.008 | 0.035 |
| rWT control<sup>c</sup> | Reverse genetics recombinant virus | 0.015 | 0.056 |
| rWA1 + E484K | Reverse genetics recombinant virus containing E484K | 0.010 | 0.047 |
| rWA1 + E484Q | Reverse genetics recombinant virus containing E484Q | 0.014 | 0.067 |
| rWA1 +L452R | Reverse genetics containing L452R | 0.008 | 0.052 |
| Omicron TVP23328 | hCoV/EHC_C192811 C.10 | 0.016 | 0.049 |

Abbreviations
**CI** = confidence interval. **IC50** = concentration inhibiting maximal activity by 50%; **IC90** = concentration inhibiting maximal activity by 90%; **nc** = no change (difference was less than ≤5-fold); **SARS-CoV-2** = severe acute respiratory syndrome coronavirus 2.
<sup>a</sup> SARS-CoV-2 S GenBank MN908947.3. <sup>b</sup> IC50 values presented are geometric mean if multiple experiments were performed and a weighted geometric mean for the Wuhan strain (n=X). 95% CI values were included for variants tested only once. <sup>c</sup> Recombinant constructed virus

**Table 3B.** LY-CoV1404 potently neutralizes SARS-CoV-2 variants

| Lineage Category | Spike Variant | VSV platform |  | Lentiviral Platform |  |
| --- | --- | --- | --- | --- | --- |
|  |  | IC<sub>50</sub><br>μg/mL<sup>a</sup> | IC<sub>90</sub><br>μg/mL<sup>a</sup> | IC<sub>50</sub><br>μg/mL | IC<sub>90</sub><br>μg/mL |
| SARS-CoV-2 WT | Genbank MN908947.3 | 0.003 | 0.024 | 0.005 | 0.021 |
|  | D614G | NA | NA | 0.003 | 0.01 |
| B.1.1.7 Related | B.1.1.7 spike | 0.001 | 0.009 | 0.004 | 0.011 |
|  | N501Y* | 0.002 | 0.007 | 0.002 | 0.009 |
|  | N501Y + del69-70 | 0.001 | 0.016 | NA | NA |
|  | del69-70 | 0.002 | 0.011 | NA | NA |
| Subpopulations of<br>B.1.1.7 | B.1.1.7 + S494P spike | 0.002 | 0.014 | NA | NA |
|  | S494P + N501Y + del69/70 | 0.005 | 0.031 | NA | NA |
|  | S494P | 0.002 | 0.034 | NA | NA |
|  | S494R | NA | NA | 0.006 | 0.037 |
|  | B.1.1.7 + E484K spike | 0.003 | 0.011 | 0.008 | 0.022 |
|  | E484K + N501Y | 0.002 | 0.012 | NA | NA |
|  | E484K* | 0.003 | 0.021 | 0.003 | 0.011 |
|  | B.1.1.7 + L452R spike | 0.002 | 0.01 | NA | NA |
|  | L452R + N501Y | <0.001 | 0.01 | NA | NA |
|  | L452R | <0.001 | 0.012 | 0.003 | 0.01 |
|  | B.1.1.7 + F490S | 0.003 | 0.013 | NA | NA |
|  | F490S + N501Y | 0.004 | 0.016 | NA | NA |
|  | F490S | 0.004 | NA | NA | NA |
|  | F490L | NA | NA | 0.005 | 0.019 |
| B.1.351 Related | B.1.351 spike | 0.002 | 0.009 | 0.003 | 0.009 |
|  | K417N + E484K + N501Y* | 0.003 | 0.021 | 0.004 | 0.013 |
|  | K417N* | 0.002 | 0.022 | 0.003 | 0.008 |
| P.1 Related | P.1 spike | 0.002 | 0.009 | 0.002 | 0.006 |
|  | K417T + E484K + N501Y | 0.002 | 0.01 | NA | NA |
|  | K417T | 0.002 | 0.011 | NA | NA |
| B.1.427/B.1.429 Related | B.1.427/B.1.429 Spike | 0.002 | 0.012 | 0.002 | 0.007 |
|  | B.1.427/B.1.429 Spike** | 0.003 | 0.015 | NA | NA |
| B.1.617 Related | B.1.617.1 Spike | 0.002 | 0.01 | NA | NA |
|  | L452R + E484Q | 0.005 | 0.027 | NA | NA |
|  | E484Q | 0.002 | 0.009 | NA | NA |
|  | B.1.617.2 spike | 0.001 | 0.007 | NA | NA |
|  | B.1.617.2 + T95I spike | 0.002 | 0.011 | NA | NA |
|  | B.1.617.2 + A222V spike | 0.002 | 0.011 | NA | NA |
|  | T478K | 0.002 | 0.01 | NA | NA |
| Other E484K-containing<br>Lineages | B.1.526 Spike | 0.002 | 0.013 | 0.002 | 0.005 |
|  | R.1 Spike | 0.003 | 0.012 | NA | NA |
| B.1.1.519 Related | B.1.1.519 spike | 0.003 | 0.013 | NA | NA |
| B.1.1.529 | B.1.1.529 Spike | 0.002 | 0.013 | NA | NA |

Table 3B. (continued)
| Lineage Category | Spike Variant | VSV platform |  | Lentiviral Platform |  |
| --- | --- | --- | --- | --- | --- |
|  |  | IC<sub>50</sub><br>μg/mL<sup>a</sup> | IC<sub>90</sub><br>μg/mL<sup>a</sup> | IC<sub>50</sub><br>μg/mL | IC<sub>90</sub><br>μg/mL |
| BA.2 | BA.2 spike | 0.004 | 0.023 | NA | NA |
| B.1.526.1 | B.1.526.1 Spike | 0.001 | 0.008 | NA | NA |
| B.1.526.2 Related | B.1.526.2 | 0.002 | 0.01 | NA | NA |
|  | S477N | 0.005 | 0.026 | NA | NA |
| C.36 Related | R346S+L452R | 0.004 | 0.018 | NA | NA |
|  | R346S | 0.001 | 0.009 | NA | NA |
| Historic Variants of Interest | B.1.258 (N439K+del69/70) | <0.001 | 0.004 | NA | NA |
|  | N439K* | 0.002 | 0.007 | 0.002 | 0.01 |
|  | B.1.1.298 (Y453F*) | 0.004 | 0.021 | 0.003 | 0.01 |
| S1/S2 Cleavage Related | Q677H | 0.006 | 0.028 | NA | NA |
|  | Q677H + D614G | 0.005 | 0.02 | NA | NA |
|  | Q677P | <0.001 | 0.007 | NA | NA |
|  | Q677P + D614G | <0.001 | 0.004 | NA | NA |
| Additional Epitope Positions of Interest | T415I | 0.002 | 0.036 | NA | NA |
|  | L452M | <0.001 | 0.006 | NA | NA |
|  | L452Q | 0.002 | 0.018 | NA | NA |
|  | L452W | <0.001 | 0.014 | NA | NA |
|  | L455M | 0.003 | 0.017 | NA | NA |
|  | F486R | NA | NA | 0.014 | 0.045 |
|  | N487R | NA | NA | 0.001 | 0.011 |

Abbreviations: D = aspartate; del = deletion; E = glutamate; G = glycine; H = histidine; I = isoleucine; IC<sub>50</sub>, IC<sub>90</sub> = concentration inhibiting maximal activity by 50% and 90%, respectively; K = lysine; L = leucine; N = asparagine; NA = not applicable; P = proline; Q = glutamine; R = arginine; S = serine; T = threonine; Y = tyrosine.
<sup>a</sup> Absolute IC estimates presented are a result of a meta-analysis of multiple replicate experiments.
\* Viral constructs for the lentiviral platform include D614G
\*\* Viral constructs for the VSV platform include D614G

**Table 3C.**
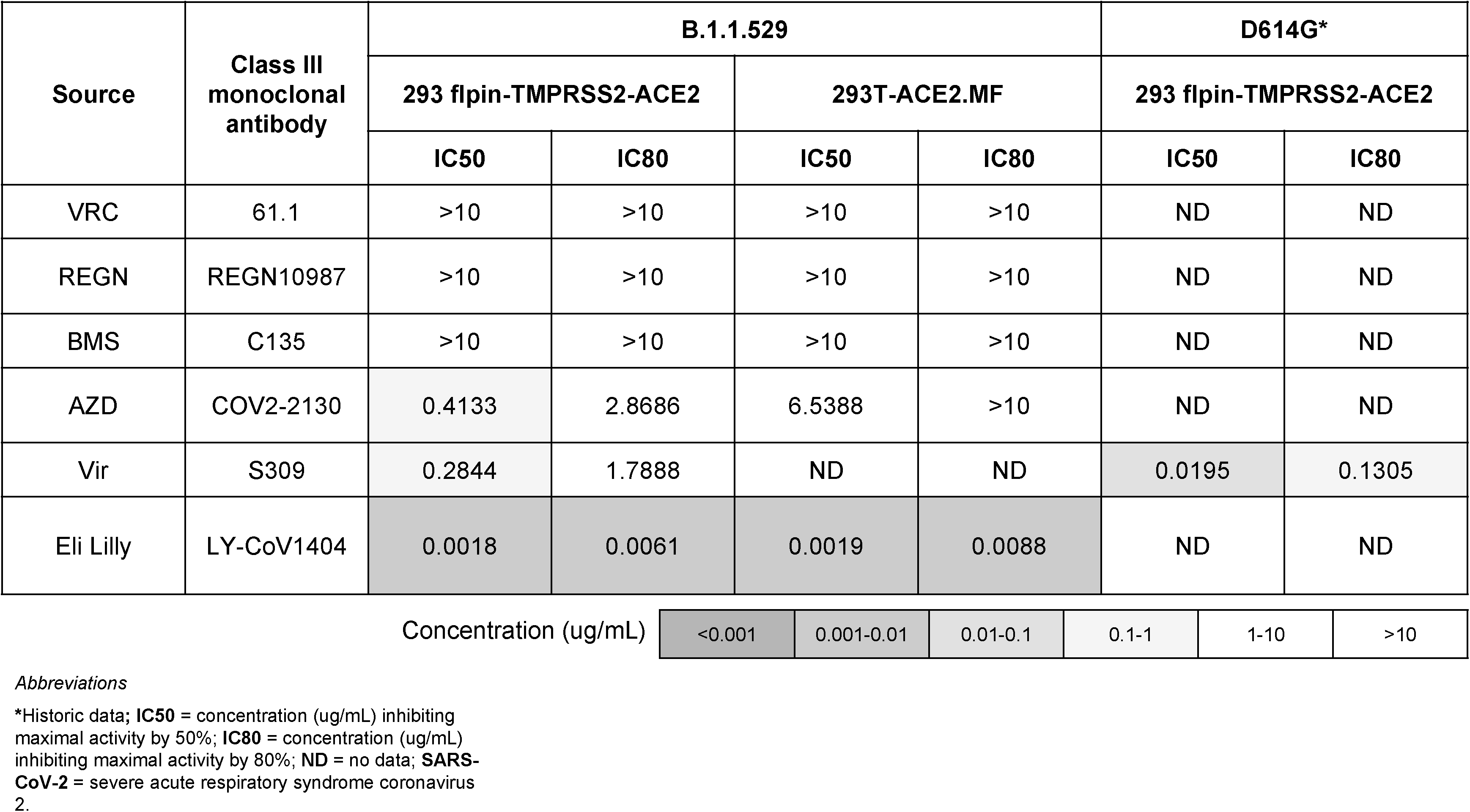
Neutralization Activity of Class III Monoclonal Antibodies Against SARS-CoV-2 on B.1.1.529

| Source | Class III monoclonal antibody | B.1.1.529 |  |  |  | D614G* |  |
| --- | --- | --- | --- | --- | --- | --- | --- |
|  |  | 293 flpin-TMPRSS2-ACE2 |  | 293T-ACE2.MF |  | 293 flpin-TMPRSS2-ACE2 |  |
|  |  | IC50 | IC80 | IC50 | IC80 | IC50 | IC80 |
| VRC | 61.1 | >10 | >10 | >10 | >10 | ND | ND |
| REGN | REGN10987 | >10 | >10 | >10 | >10 | ND | ND |
| BMS | C135 | >10 | >10 | >10 | >10 | ND | ND |
| AZD | COV2-2130 | 0.4133 | 2.8686 | 6.5388 | >10 | ND | ND |
| Vir | S309 | 0.2844 | 1.7888 | ND | ND | 0.0195 | 0.1305 |
| Eli Lilly | LY-CoV1404 | 0.0018 | 0.0061 | 0.0019 | 0.0088 | ND | ND |

|  |  |  |  |  |  |  |
| --- | --- | --- | --- | --- | --- | --- |
| Concentration (ug/mL) | <0.001 | 0.001-0.01 | 0.01-0.1 | 0.1-1 | 1-10 | >10 |

*Abbreviations* \*Historic data; **IC50** = concentration (ug/mL) inhibiting maximal activity by 50%; **IC80** = concentration (ug/mL) inhibiting maximal activity by 80%; **ND** = no data; **SARS-CoV-2** = severe acute respiratory syndrome coronavirus 2.

**Table 3D.**
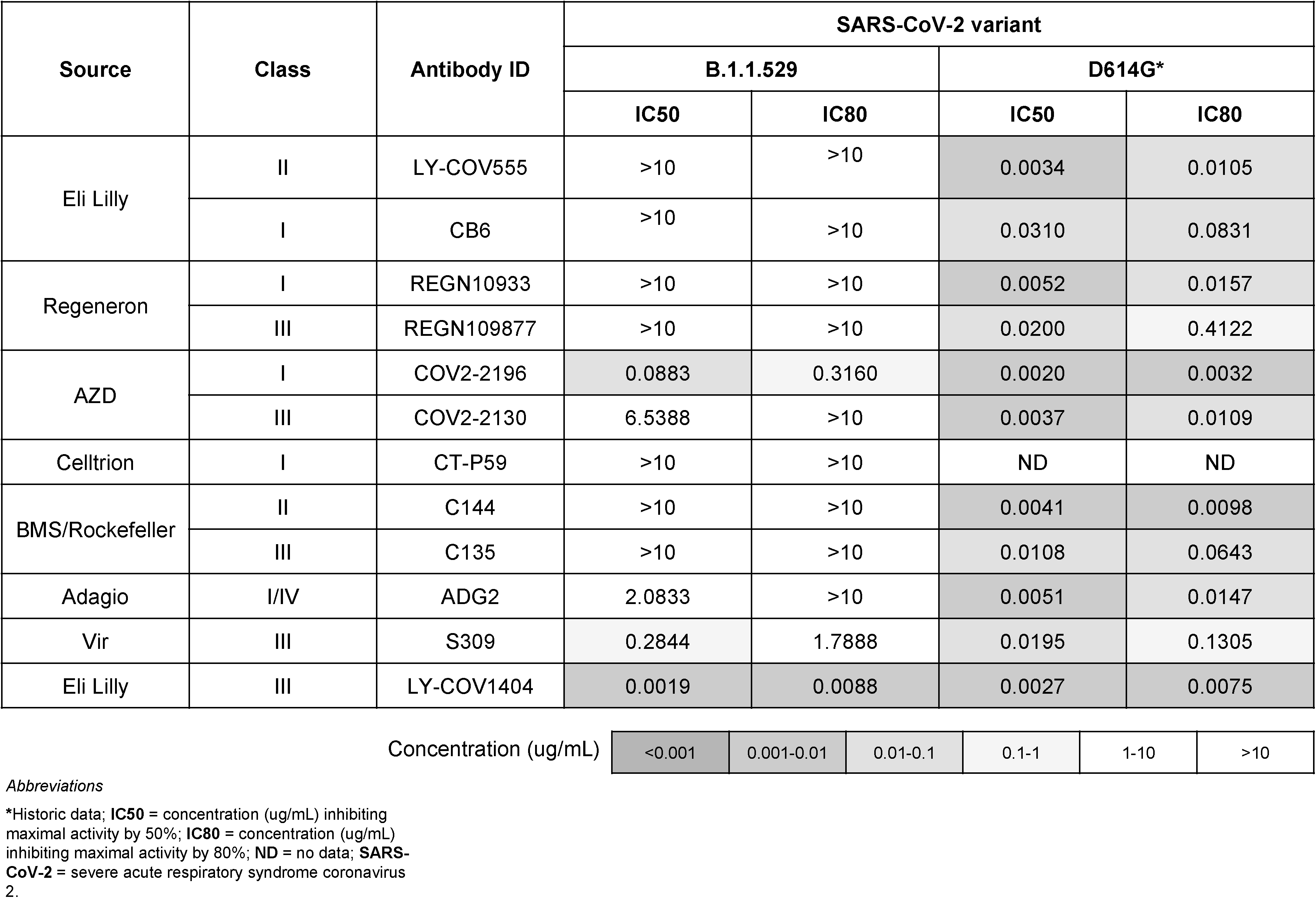
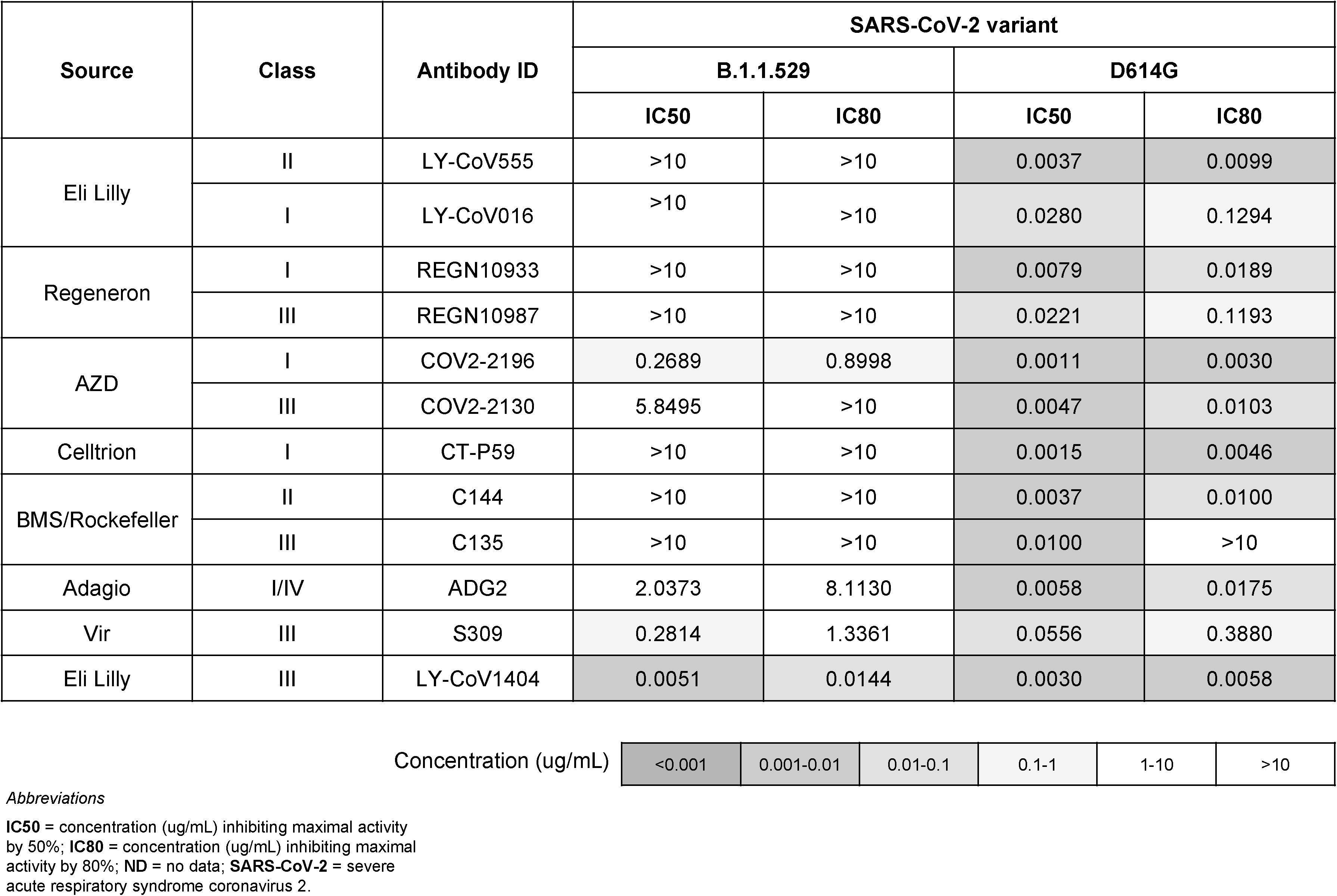
Neutralization Activity of Clinically-Relevant Monoclonal Antibodies Against the SARS-CoV-2 variants B.1.1.529 and D614G

### LY-CoV1404 blocks ACE2-Spike interaction and binds to an infrequently mutated RBD epitope

To fully characterize the binding epitope of LY-CoV1404 we determined the X-ray crystal structure of LY-CoV1404 Fab bound to S protein RBD and CryoEM structure of intact S protein bound to LY-CoV1404 (**Figure 3**). The Fab of LY-CoV1404 binds to a region overlapping the ACE2-interacting site of the S protein that is accessible in both the “up” and “down” conformers of the RBD on the S protein (**Figure S2**). While this property would suggest that LY-CoV1404 is a Class 2 binder (Barnes et al., 2020), the structural location of the epitope is closer to the canonical Class 3 binder, VIR-S309 (Wang et al., 2021b). Interestingly, the binding epitope of LY-CoV1404 is very similar to previously described antibodies imdevimab (REGN10987) (Hansen et al., 2020) and Fab 2-7 (Cerutti et al., 2021; Liu et al., 2020). This similarity is apparent from the structural superposition of the Fabs as well as which RBD residues interact with the antibodies (**Figure S3**). Furthermore, though independently discovered, LY-CoV1404 and Fab 2-7 (Cerutti et al., 2021) share 92% amino acid sequence identity in the variable regions of both their heavy and light chains, and they both engage the RBD similarly through their heavy and light chains. By contrast, REGN10987 is more sequence divergent and nearly all of its interactions with the RBD are through its heavy chain. LY-CoV1404 has a contact surface area on the RBD of 584 Å^2^, compared to 496 Å^2^ for Fab 2-7, and only 343 Å^2^ for REGN10987 (**Table S2**).

**Figure 3.**
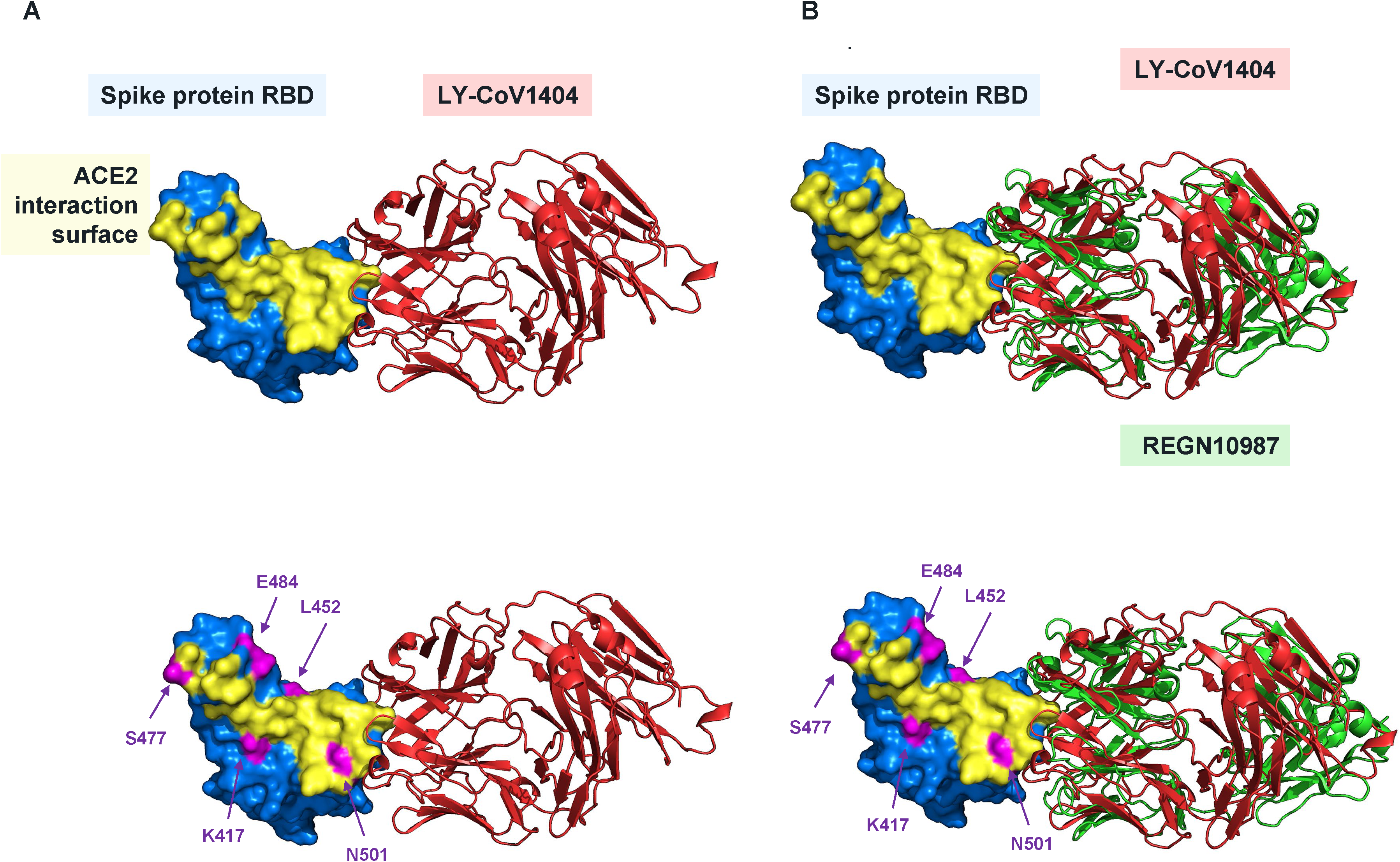
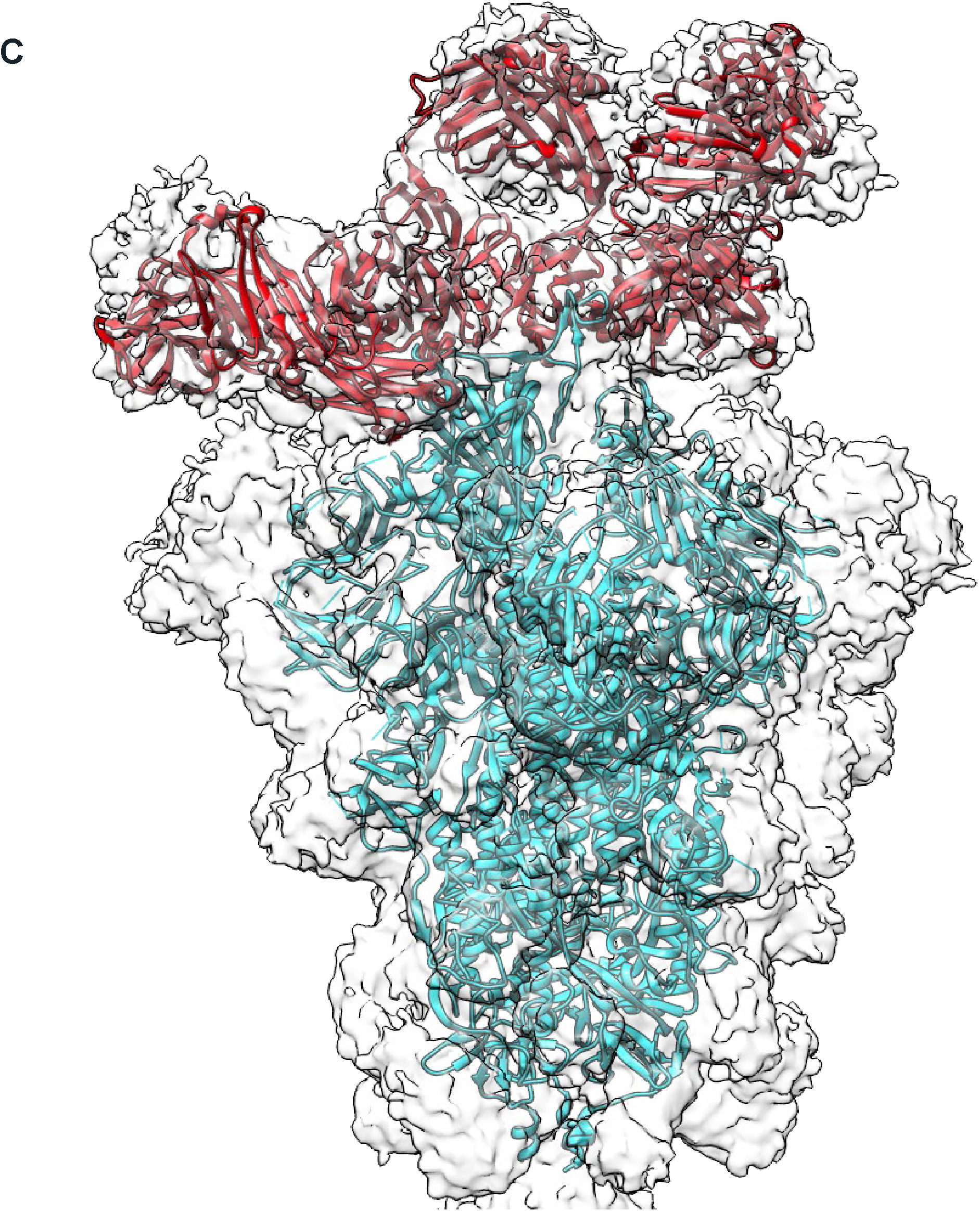
Structural analysis of LY-CoV1404 binding to RBD. (A) Three-dimensional structure of the Fab portion of LY-CoV1404 bound to the spike protein receptor-binding domain. (B) Overlay of LY-CoV1404 compared to imdevimab (REGN10987, PDB 6DXG), bound to the spike protein receptor-binding domain. (C) CryoEM density map of S protein-LY-CoV1404 complex shown at low threshold and docked with atomic model of S protein (cyan) and LY-CoV1404 (red) highlights binding of 3 fab molecules in 1 RBD “up” and 2 “down” conformation.

To determine the frequency of mutations of the amino acid residues of the S protein that are close contacts with LY-CoV1404 as determined by structural analysis, we used the publicly deposited mutations observed at these sites in the GISAID EpiCoV database (**Table 4**). There were very few changes to these identified amino acids, with the only two residues below the >99% unchanged threshold: N439 and N501 (reported as having 99.371% and 78.189% conservation at these positions respectively), demonstrating that the LY-CoV1404 epitope has remained relatively unchanged during the pandemic. We identified the most prevalent mutations at epitope residues in the GISAID EpiCoV database and characterized their impact on the functional activity of LY-CoV1404. Interestingly, the observed N439K and N501Y mutations had no impact on LY-CoV1404 binding or neutralization (**Table 5**). The lack of effect by the N493K mutation is interesting in the context of the loss of antibody binding previously observed for REGN10987 (Thomson et al., 2021) demonstrating that beyond the epitope, the specific interactions are critical to determining whether resistant mutations will arise for any particular antibody. A loss of binding, ACE2 competition and neutralization activity was observed with mutations at positions 444 and 445 (which could arise by a single nucleotide change) (**Table 5**). Importantly however, these changes are extremely rare in the general population as reported in the GISAID database (**Table 4**). The ability of LY-CoV1404 to maintain neutralization potency despite a variety of mutations that have been shown to nullify activity of several other potent neutralizing antibodies (Rappazzo et al., 2021) indicates that this antibody may be uniquely suited to combat the current VOC. Critically, potent activity against all of these variants suggests that LY-CoV1404 binds to an epitope that has acquired few mutations.

**Table 4.** Summary of LY-CoV1404-Binding Epitope Positions and their Conservation in GISAID (as of 12/21/20201)

| RBD Contact Residue | % Conservation in GISAID<sup>a</sup> |
| --- | --- |
| T345 | 99.988 |
| R346 | 99.652 |
| N439 | 97.371 |
| N440 | 99.742 |
| L441 | 99.992 |
| S443 | 99.999 |
| K444 | 99.964 |
| V445 | 99.986 |
| G446 | 99.744 |
| G447 | 99.997 |
| N448 | 99.996 |
| Y449 | 99.955 |
| N450 | 99.968 |
| Q498 | 99.740 |
| P499 | 99.979 |
| T500 | 99.997 |
| N501 | 78.189 |
| G502 | 99.999 |
| V503 | 99.977 |
| Q506 | 99.999 |
| R509 | 99.999 |

*Abbreviations*
**GISAID** = Global Initiative on Sharing All Influenza Data; **RBD** = receptor binding domain.
<sup>a</sup> % Conservation is defined as the percentage of deposited sequences containing the nonmutated amino acid in GISAID (as of February 2021).

**Table 5.**
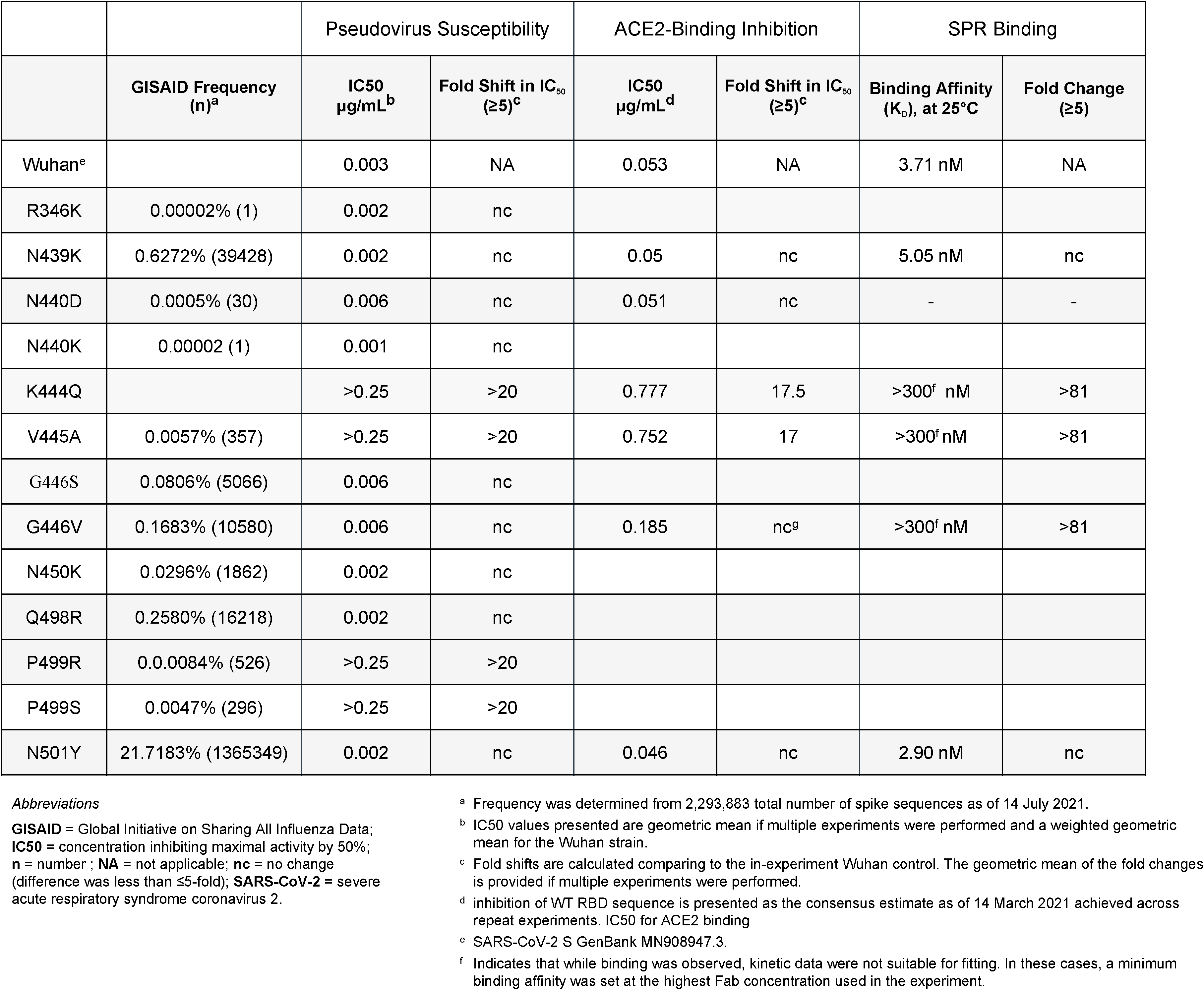
LY-CoV1404 Activity Against SARS-CoV-2 Variant Proteins and Viruses Located in the LY-CoV1404-Binding Site

With the exception of N439 and N501, the residues with which LY-CoV1404 interacts are not prevalent in current VOC (e.g. 417, 439, 452, 484). Importantly, while the LY-CoV1404 binding epitope includes residues both N439 and N501, LY-CoV1404 still binds the N501Y mutant (**Figure 1F**), and variants B.1.1.7 and B.1.351, which both carry the N501Y mutation and are neutralized as potently as wild type virus in both the pseudovirus and live virus assays (**Table 3A/B**). Similarly, the LY-CoV1404 binding epitope includes residue N439, and substantially overlaps with the binding site of the clinically validated antibody imdevimab (**Figure 3B**, **Figure S2**). However, unlike imdevimab (Thomson et al., 2021), LY-CoV1404 retains full functional neutralization against pseudovirus with the N439K mutant (**Table 5**), indicating that these residues do not play critical roles in LY-CoV1404 interaction with S protein. In addition, the mutations present in B.1.1.529 that reside within the binding epitope for LY-CoV1404, specifically G446S, N440K, Q498R, and N501Y, do not impact the neutralization activity of LY-CoV1404. Importantly, potent activity against all the tested variants suggests LY-CoV1404 binds uniquely to an epitope that has acquired few mutations and is not sensitive to the mutations that have arisen to date.

### Mutational analysis of S protein binding to LY-CoV1404

We explored the potential for resistant mutations to arise by yeast-display of the S protein RBD. A library was constructed in which all amino acid residues were sampled at positions 331 to 362 and 403 to 515 of the spike glycoprotein containing the epitope of LY-Cov1404. Cells containing RBD substitutions that were still able to bind soluble ACE2 after antibody treatment were isolated and sequenced to determine the location and identity of the change. Further functional characterizations of isolates were limited to those containing a change of a single amino acid residue which can be generated by a single nucleotide change based on the codon used in the Wuhan isolate sequence.

Consistent with the structure of the RBD complex, selection using LY-CoV1404 identified potential susceptibility to some, but not all, substitutions at residues K444, V445, G446, and P499. None of the observed G446 changes identified from the selection could arise from a single nucleotide substitution but nevertheless suggested a potentially liable residue with LY-CoV1404. The presence of the G446V variant in the GISAID database, as well as the ability of this variant to be resistant to imdevimab treatment prompted further analysis. The ability of LY-CoV1404 to inhibit ACE2 binding to RBD substitutions that have been identified in common circulating viral variants and/or have been shown to impart resistance to the authorized antibodies bamlanivimab, etesevimab, casirivimab, and imdevimab was also analyzed (**Table 6**).

**Table 6.**
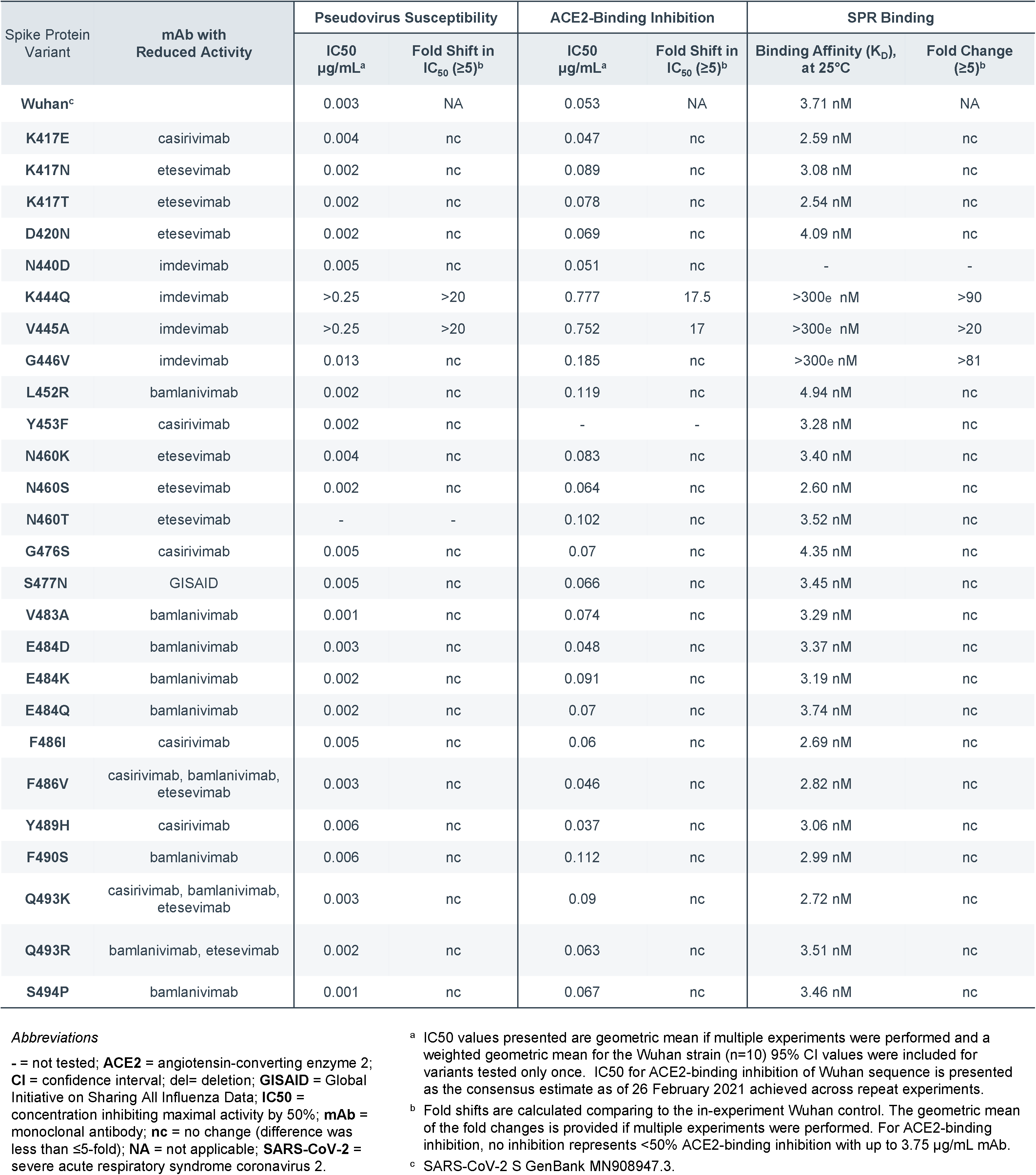
LY-CoV1404 Activity against SARS-CoV-2 Variant Viruses with Known Resistance to Other Clinical Monoclonal Antibodies

| Spike Protein Variant | mAb with Reduced Activity | Pseudovirus Susceptibility |  | ACE2-Binding Inhibition |  | SPR Binding |  |
| --- | --- | --- | --- | --- | --- | --- | --- |
|  |  | IC50 µg/mL<sup>a</sup> | Fold Shift in IC<sub>50</sub> (≥5)<sup>b</sup> | IC50 µg/mL<sup>a</sup> | Fold Shift in IC<sub>50</sub> (≥5)<sup>b</sup> | Binding Affinity (K<sub>D</sub>), at 25°C | Fold Change (≥5)<sup>b</sup> |
| Wuhan<sup>c</sup> |  | 0.003 | NA | 0.053 | NA | 3.71 nM | NA |
| K417E | casirivimab | 0.004 | nc | 0.047 | nc | 2.59 nM | nc |
| K417N | etesevimab | 0.002 | nc | 0.089 | nc | 3.08 nM | nc |
| K417T | etesevimab | 0.002 | nc | 0.078 | nc | 2.54 nM | nc |
| D420N | etesevimab | 0.002 | nc | 0.069 | nc | 4.09 nM | nc |
| N440D | imdevimab | 0.005 | nc | 0.051 | nc | - | - |
| K444Q | imdevimab | >0.25 | >20 | 0.777 | 17.5 | >300<sub>e</sub> nM | >90 |
| V445A | imdevimab | >0.25 | >20 | 0.752 | 17 | >300<sub>e</sub> nM | >20 |
| G446V | imdevimab | 0.013 | nc | 0.185 | nc | >300<sub>e</sub> nM | >81 |
| L452R | bamlanivimab | 0.002 | nc | 0.119 | nc | 4.94 nM | nc |
| Y453F | casirivimab | 0.002 | nc | - | - | 3.28 nM | nc |
| N460K | etesevimab | 0.004 | nc | 0.083 | nc | 3.40 nM | nc |
| N460S | etesevimab | 0.002 | nc | 0.064 | nc | 2.60 nM | nc |
| N460T | etesevimab | - | - | 0.102 | nc | 3.52 nM | nc |
| G476S | casirivimab | 0.005 | nc | 0.07 | nc | 4.35 nM | nc |
| S477N | GISAID | 0.005 | nc | 0.066 | nc | 3.45 nM | nc |
| V483A | bamlanivimab | 0.001 | nc | 0.074 | nc | 3.29 nM | nc |
| E484D | bamlanivimab | 0.003 | nc | 0.048 | nc | 3.37 nM | nc |
| E484K | bamlanivimab | 0.002 | nc | 0.091 | nc | 3.19 nM | nc |
| E484Q | bamlanivimab | 0.002 | nc | 0.07 | nc | 3.74 nM | nc |
| F486I | casirivimab | 0.005 | nc | 0.06 | nc | 2.69 nM | nc |
| F486V | casirivimab, bamlanivimab, etesevimab | 0.003 | nc | 0.046 | nc | 2.82 nM | nc |
| Y489H | casirivimab | 0.006 | nc | 0.037 | nc | 3.06 nM | nc |
| F490S | bamlanivimab | 0.006 | nc | 0.112 | nc | 2.99 nM | nc |
| Q493K | casirivimab, bamlanivimab, etesevimab | 0.003 | nc | 0.09 | nc | 2.72 nM | nc |
| Q493R | bamlanivimab, etesevimab | 0.002 | nc | 0.063 | nc | 3.51 nM | nc |
| S494P | bamlanivimab | 0.001 | nc | 0.067 | nc | 3.46 nM | nc |

*Abbreviations*
- = not tested; **ACE2** = angiotensin-converting enzyme 2; **CI** = confidence interval; del= deletion; **GISAID** = Global Initiative on Sharing All Influenza Data; **IC50** = concentration inhibiting maximal activity by 50%; **mAb** = monoclonal antibody; **nc** = no change (difference was less than ≤5-fold); **NA** = not applicable; **SARS-CoV-2** = severe acute respiratory syndrome coronavirus 2.
<sup>a</sup> IC50 values presented are geometric mean if multiple experiments were performed and a weighted geometric mean for the Wuhan strain (n=10) 95% CI values were included for variants tested only once. IC50 for ACE2-binding inhibition of Wuhan sequence is presented as the consensus estimate as of 26 February 2021 achieved across repeat experiments.
<sup>b</sup> Fold shifts are calculated comparing to the in-experiment Wuhan control. The geometric mean of the fold changes is provided if multiple experiments were performed. For ACE2-binding inhibition, no inhibition represents <50% ACE2-binding inhibition with up to 3.75 µg/mL mAb.
<sup>c</sup> SARS-CoV-2 S GenBank MN908947.3.

We further characterized LY-CoV1404 activity in the presence of mutations known to weaken the effectiveness of other antibodies in clinical testing. Of the mutations tested, only K444Q, V445A, and G446V impacted LY-CoV1404 binding (**Table 5**). Interestingly, while the K444Q and V445A mutations also resulted in reduced ACE2 competition and neutralization activity, the G446V mutation only appeared to significantly impact the monomeric binding affinity, along with modest changes in ACE2 competition and neutralization. These residues are very rarely mutated according to GISAID reports (**Tables 4**,**5**), thus suggesting a low risk of mutations emerging at these residues and restriction in functional activity of LY-CoV1404. We examined the frequency of all variants observed at residues 444 and 445 within the GISAID-EpiCov database (Shu and McCauley, 2017), as of December 21, 2021; see Materials and Methods).

These two sites were rarely mutated, with 0.036% at residue 444 and 0.014% at residue 445 of samples collected contain mutations. It is not known whether all variants at these locations would confer resistance. Of the specific mutations tested (**Table 5**), we find that only 0.0057% of samples contained the known resistance V445A, while no occurrences of K444Q were observed. This suggests LY-CoV1404 may remain effective even as new strains emerge. These data also indicate that LY-CoV1404 is not only likely to be highly effective against current variants, but is less likely to be impacted by future mutations given the low level of changes observed to date in its binding epitope (McCormick et al., 2021).

## Discussion

We report the discovery of LY-CoV1404, a highly potent SARS-CoV-2 S protein RBD-binding antibody that maintains binding and neutralizing activity across currently known and reported VOC. LY-CoV1404 binds to or neutralizes variants including B.1.1.529 (Omicron) and those first identified in India (B.1.617.1, B.1.617.2, and B.1.617.3), the UK (B.1.1.7), South Africa (B.1.351), Brazil (P.1), California (B.1.426 and B.1.429), and New York (B.1.526). The neutralizing potency of LY-CoV1404 against either pseudotyped viral mutant reporters or authentic virus indicates that the antibody binds to an epitope on the S protein that is accessible and bound with high affinity by LY-CoV1404. In addition, LY-CoV1404 blocks interaction between ACE2 and the S protein, providing a strong, well-documented mechanism for the potent neutralizing activity. Multiple antibodies that rely upon this receptor-blocking mechanism have recently been developed as treatments for COVID-19 and clearly demonstrate that antibody-mediated viral neutralization is a safe and highly efficacious COVID-19 treatment strategy (Chen et al., 2020; Eli Lilly and Company, 2020; Gottlieb et al., 2021; Regeneron, 2021; Tada, et al., 2021). However, the emergence of SARS-CoV-2 variants has diminished the effectiveness of several of these therapeutic antibodies (Davies et al.; Munnink et al., 2021; Plante et al., 2020; Tegally et al., 2021). The rapid spread of a new VOC (B1.1.529, Omicron) has diminished the function of all the authorized and clinically tested antibodies. Notably, this is true even for two antibodies that have demonstrated cross-reactivity to SARS and were believed to be less susceptible to viral escape; ADG20 was found to be more than 300-fold less potent against B1.1.529, while the potency of S309 was reduced by approximately 10-fold. This underscores the unpredictability of SARS-CoV-2 spike mutation. Importantly, LY-CoV-1404 is the only clinically tested antibody that retains full functional potency in pseudotype virus neutralization assays against this mutant. Our data suggest that LY-CoV-1404 is more than 50-fold more potent against omicron than any of the other clinical stage antibodies we have tested. In addition, either Omicron or the BA.2 subvariant of Omicron have been reported to severely reduce or eliminate neutralization activity against all clinically tested antibodies (Iketani et al., 2022). LY-CoV1404 retains full potency against both variants (Fig 3B) in pseudovirus assays. The unique binding epitope of LY-CoV1404, together with the low frequency of mutations observed within this epitope, indicate that this antibody could provide an effective therapeutic option against current VOCs and emerging variants as a complementary approach to vaccinations and other COVID-19 therapies.

With more than 404 million infections world-wide it is not surprising that SARS-CoV-2 variants with a potential selective advantage have arisen. These resulting variants have acquired either increased affinity for ACE2 binding (Altmann et al., 2021), increased transmissibility and infection severity (Gu et al., 2020), or increased immune evasion (Altmann et al., 2021), and threaten to reduce the effectiveness of currently available vaccinations (Mansbach et al., 2020; Yurkovetskiy et al., 2020). The D614G mutation, reported early in the course of the pandemic, has been determined to stabilize the S protein, leading to more efficient ACE2 interactions and infections (Santos and Passos, 2021). The rapid spread of the B.1.1.7 variant, which contains the key mutation N501Y, has been correlated with increased ACE2 binding affinity (Baric, 2020; Horby et al., 2021; Plante et al., 2020). Certain recurring mutations within the S protein have also been reported in variants from different geographical regions indicating that these common mutations may confer an advantage to the virus (Kuzmina et al., 2021; Liu et al., 2021; Rees-Spear et al., 2021). Reports of particular key mutations such as E484K are present in several variants emerging independently in several geographic locations (South Africa B.1.351, Brazil P.1, New York B.1.526, and recently some UK B.1.1.7 strains) (Xie et al., 2021). Investigations have indicated that vaccine-induced and natural immune responses may not be as effective against variants carrying the E484K mutation (Plante et al., 2020). In addition, several other specific mutations within the S protein have been demonstrated to reduce binding and effectiveness of antibody therapies (Eli Lilly and Company, 2020; Regeneron, 2021). With the proven clinical effectiveness of antibody therapies at preventing hospitalization, reducing severity of disease symptoms and death (Darby and Hiscox, 2021), the importance of access to effective antibody treatments targeting conserved neutralizing epitopes for response to viral variants must be recognized.

It remains unknown whether vaccines will significantly alter the mutation profile of the virus (Choi et al., 2020; Kemp et al., 2021). If, as vaccination continues world-wide, the virus responds to this selection pressure with particular mutations to avoid the effects of vaccination, it will be important to maintain alternative therapies to help overcome these variants. It has been hypothesized that certain variants may have arisen in immunocompromised individuals, which provided the virus with an extended timeframe for continued viral replication and the accumulation of many mutations, seeding these novel VOC (Aschwanden, 2021). A potent neutralizing monoclonal antibody, such as LY-CoV1404, would be a potential therapeutic option for these immunocompromised patients, with rapid neutralization of the virus providing both protection for the patient, and less opportunity for viral mutation and evolution. Finally, immunocompromised individuals are less likely to respond well to vaccines, further highlighting the utility of monoclonal antibody therapy for protecting this population. Monoclonal antibodies with potent neutralizing effects could provide a safe and robust medical countermeasure against variants or a necessary, complementary alternative form of protection for those individuals who cannot, or do not receive vaccines. For the reasons listed above, it has been proposed that achieving “herd immunity” is unlikely in the near term for SARS-CoV-2 (Cathcart et al., 2021; Pinto et al., 2020), further emphasizing the need for medical alternatives to combat viral resistance.

Recently, an antibody that binds to an epitope of the SARS-CoV-2 S protein that is distinct from current VOCs has been described. This antibody, VIR-7831(GSK, 2021), binds to and neutralizes current variants of concern, including Delta (B.1.617.2) and Omicron (B.1.1.529), and has recently been reported to have clinical efficacy at a 500mg dose (Cathcart et al., 2021). LY-CoV1404 is equally effective in viral neutralization against all tested variants and importantly is several-fold more potent in viral neutralization assays (Cathcart et al., 2021) (**Figure 2B**). Though it remains to be tested, this increased potency has the potential to enable lower doses and subcutaneous administration for either treatment or prophylaxis. Interestingly, VIR-7831 was derived from the antibody S309, which was identified from a SARS-CoV-1 convalescent patient (Cathcart et al., 2021). This antibody is cross-reactive to SARS and binds to one of the few epitopes shared between the viruses, which includes an N-linked glycan. By comparison, LY-CoV1404 was discovered from a SARS-CoV-2 convalescent patient and selected based on its cross-reactivity to SARS-CoV-2 variants, thereby allowing testing and identification of much more potent neutralizing epitopes. A more recently identified antibody from a COVID-19-recovered patient, S2X259, was described with broad cross-reactivity to variants (including B.1.1.7, B.1.351, B.1.429, and P.1) and several zoonotic strains (Tortorici et al., 2021). This antibody, while able to bind widely to coronaviruses, had neutralizing capacity against SARS-CoV-2 in authentic virus-neutralization assays that was substantially less potent than LY-CoV1404. Finally, an engineered antibody with broad cross-reactivity against coronaviruses and potent neutralization was recently described (Rappazzo et al., 2021), which was optimized for binding following identification from a SARS convalescent patient.

Mutations within the S protein and in particular in the RBD are inevitable, particularly as SARS-CoV-2 has been present in the human population for only 18 months (Jones et al., 2020). Since the pandemic began, we have continued to screen patient samples and have identified thousands of human antibodies with associated functional data. We analyzed this antibody database in response to the increase in VOC and identified many potential solutions based on these variants. LY-CoV1404 is one example of this screening and analysis. In order to fully prepare for the inevitability of additional mutants and reduction in treatment effectiveness, we propose expanding this effort into a panel of potent neutralizing antibodies with full coverage of the S protein RBD. The antibodies in this panel, if well characterized and advanced to (at minimum) research cell banks (RCBs), could accelerate key manufacturing steps and be rapidly deployed. This strategy of a panel of antibodies would enable a rapid response to any emerging variant, necessitating only mutation identification and assessment of binding and neutralization of a new mutation arising in a particular geography. The rapid deployment of the antibody into clinical testing could follow immediately after identification of novel VOC. This stands as a desirable rapid countermeasure to safeguard ongoing pandemic responses, especially as antibodies, if administered early in the course of disease or prophylactically, have demonstrated to be safe and effective for combatting SARS-CoV-2 infection. The antibody described in this report, LY-CoV1404, is a key member of such a panel, especially given its retention of potency against a variety of emerging viral variants. LY-CoV1404 received Emergency Use Authorization on February 11, 2022.

### Limitations of the study

Our work focused on the capacity of the described antibody to bind to and neutralize SARS-CoV-2 by interaction with the spike protein. The neutralization of SARS-CoV-2 by LY-CoV1404 and other neutralizing antibodies was measured *in vitro*. Further studies using *in vivo* models of COVID-19 infection will help elucidate the mechanisms of LY-CoV1404 in treating and/or preventing disease.

## Author contributions

B.E.J., J.D., C.C., B.A.H., D.F., J.H., V.d.P., M.W., E.L., M.R., L.D., A.O.N., R.v.d.L, P.P, H.D., F.A.G., I.L., L.A., C.P., K.D., D.K., J.A., B.S.G., J.R.M, N.K., J.J.F., S.H., I.H., L.M., H.C.P., B.R., and R.E.H. conceived of and designed experiments, data analysis and reporting, and participated in manuscript authoring and review. A.P. and J.H. conceived of and designed experiments (crystallography/structure determination) and participated in manuscript authoring and review. K.S.C., P.V., L.W., E.S.Y., Y.Z., T.Z., J.M., P.D.K., N.J.S., and W.S. conceived of, designed and performed experiments/reagents (pseudovirus neutralization assay), data analysis and reporting, and participated in manuscript authoring and review. R.S. and J.C. performed yeast-display experiments to identify resistance mutations and characterized ACE2 competition for variants. T.W.G., R.W.C., D.K., and J.A. conceived of and designed experiments (PRNT assay), data analysis and reporting, and participated in manuscript review. C.C., J.D., K.E.H, and C.V.B conceived of and designed experiments (IFA assay), data analysis and reporting, and participated in manuscript review. R.G. participated in conception and design of bioinformatics experiments. data analysis and reporting. M.A.S. designed and implemented software improvements to Celium^TM^ for antibody selection. R.G., M.A.S., K.J. and D.W.C., assisted in the acquisition, organization and interpretability of the data, and participated in manuscript review. S.Ž. and K.W. participated in design, execution, data analysis and interpretation (screening and validation experiments), as well as drafting and review of this manuscript. K.W. participated in the interpretation of screening, validation and characterization data for downselection of antibodies for expression and characterization. L.K. and Y.H. designed and executed binding kinetics & epitope binning experiments, data analysis and reporting, and participated in manuscript authoring and review. D.P. designed and implemented data analysis and reporting pipelines for binding kinetics, epitope binning and ACE2 blocking, and participated in manuscript authoring and review. P.X. conceived experiments, designed and established the fast mutant full-length spike proteins expression vector generation protocol for cell-based validation, performed mutant tracing and surveys, and participated in manuscript authorship. P.S. participated in design of data acquisition pipeline (discovery, characterization, bioinformatics and antibody lead selection); T.L.F participated in data analysis and reporting, manuscript authorship and review. B.C.B., C.L.H. and E.F. conceived and designed experiments, analyzed and reported data (discovery, characterization, bioinformatics and antibody lead selection) and participated in manuscript authorship and review.

## Acknowledgments

We would like to thank the following: Kristi Huntington (Eli Lilly and Company) and Emma McCarren (AbCellera Biologics Inc.) for project leadership and coordination; Franz Trianna, Douglas Burtrum, Prabakaran Narayanasamy, Xiaomin Yang, Ricky Lieu, Dongmei He, Henry Koo, of Eli Lilly and Company for reagent and antibody cloning, expression and purification; Craig Dickinson, Kristina Coleman and Jeffrey Boyles of Eli Lilly and Company for initiation and reagent generation for crystallography experiments; David Driver of Eli Lilly & Company for assistance with pseudovirus genomic titering; Natalie Thornburg, Kenneth Plante, and the University of Texas Medical Branch (UTMB) World Reference Collection for Emerging Viruses and Arboviruses (WRCEVA) for the WA-1 isolate (deposited by N. Thornburg), San Diego isolate (deposited by N. Thornburg), the Maryland isolate (Mehul Suthar), and the Delta isolate; the SARS-CoV-2/INMI-1-Isolate/2020/Italy used in this publication was kindly provided by the European Virus Archive Goes Global project that has received funding from the European Union’s Horizon 2020 research and innovation program under grant agreement no. 653316 Steven Widen of the NextGen Sequencing Core at UTMB for genomic sequencing of the variants used in this study; the SARS-CoV-2/INMI-1-Isolate/2020/Italy used in this publication was kindly provided by the European Virus Archive Goes Global project that has received funding from the European Union’s Horizon 2020 research and innovation program under grant agreement no. 653316; Sherie Duncan, Anders Klaus, Keith Mewis, Karine Herve, Amanda Moreira, and Emilie Lameignere of AbCellera Biologics Inc. for technical support; Chad Thiessen of AbCellera Biologics Inc. for development of features for Celium^TM^ required for antibody selection; Clara Ng-Cummings of AbCellera Biologics Inc. for figure generation; Wolfgang Glaesner of Eli Lilly and Company for management and personnel resources. We gratefully acknowledge the authors from the originating laboratories and the submitting laboratories, who generated and shared via GISAID genetic sequence data, on which this research is based.

## Disclaimer

This research was, in part, funded by the U.S. Government. The views and conclusions contained in this document are those of the authors and should not be interpreted as representing the official policies, either expressed or implied, of the U.S. Government.

Opinions, conclusions, interpretations, and recommendations are those of the authors and are not necessarily endorsed by the U.S. Army. The mention of trade names or commercial products does not constitute endorsement or recommendation for use by the Department of the Army or the Department of Defense.

This research used resources of the Advanced Photon Source, a U.S. Department of Energy (DOE) Office of Science User Facility operated for the DOE Office of Science by Argonne National Laboratory under Contract No. DE-AC02-06CH11357. https://www.aps.anl.gov/Science/Publications/Acknowledgment-Statement-for-Publications. Use of the Lilly Research Laboratories Collaborative Access Team (LRL-CAT) beamline at Sector 31 of the Advanced Photon Source was provided by Eli Lilly & Company, which operates the facility. http://lrlcat.lilly.com/

## STAR Methods

### Lead contact

Further information and requests for resources and reagents should be directed to and will be fulfilled by the Lead Contact, Dr. Bryan E. Jones.

### Materials availability

SARS-CoV-2 viruses are being made available through the Biodefense and Emerging Infections Research Resources Repository. Antibodies for non-commercial internal research purposes can be obtained from AbCellera Biologics Inc. and/or Eli Lilly and Company under material transfer agreements (MTA) . Purified proteins for *in vitro* experiments can be generated upon execution of a material transfer agreement (MTA) with inquiries directed to Dr. Bryan E. Jones.

### Data and code availability

Atomic coordinates of the X-ray crystal structure of the LY-CoV1404:RBD complex have been deposited in the Protein Data Bank under accession codes 7MMO. This work is licensed under a Creative Commons Attribution 4.0 International (CC BY 4.0) license, which permits unrestricted use, distribution, and reproduction in any medium, provided the original work is properly cited. To view a copy of this license, visit https://creativecommons.org/licenses/by/4.0/. This license does not apply to figures/photos/artwork or other content included in the article that is credited to a third party; obtain authorization from the rights holder before using this material.

This paper does not report original code.

Any additional information required to reanalyze the data reported in this paper is available from the lead contact upon request.

### Data

All data associated with this study are available in the main text or the supplementary materials and will be shared by the lead contact upon request.

### Experimental model details

#### Cells and viruses

Chinese hamster ovary (CHO) cells and human embryonic kidney (HEK-293) were obtained from ATCC and cultured in Dulbecco’s modified Eagle medium (DMEM) supplemented with 1% penicillin-streptomycin (ThermoFisher) and 10% FBS (ThermoFisher).

PBMC samples were collected under institutional review board (IRB)-approved protocols as part of the Hospitalized and Ambulatory Adults with Respiratory Viral Infections (HAARVI) study at the University of Washington (protocol #STUDY00000959) and Vaccine Research Center (VRC), National Institute of Allergy and Infectious Diseases (NIAID) and National Institutes of Health (NIH; protocol-VRC400, NIH-07IN194). Cells were thawed, activated in culture to generate antibody secreting cells and enriched for B cell lineage cells prior to injection into AbCellera’s microfluidic screening devices with either 91,000 or 153,000 individual nanoliter-volume reaction chambers (U.S Patent 10421936B2).

Live cell assays used passively dyed suspension-adapted CHO cells transiently transfected to surface-express full-length SARS-CoV-2 spike protein (GenBank ID MN908947.3) with a green fluorescent protein (GFP) reporter, and non-transfected cells as a negative control.

Vero-E6 cells were maintained in plaque assay media (complete MEM media with 1% BGS + 1% low melting point agarose).

Authentic SARS-CoV-2 virus (MT020880.1) was obtained through the Biodefense and Emerging Infections Research Resources Repository.

#### Recombinant proteins

Recombinant antibodies and other proteins were produced as previously described (Jones et al., 2020). The antigen binding fragment (Fab) portion of LY-CoV1404 was generated by proteolytic digestion using immobilized papain (ThermoFisher Scientific), followed by removal of un-cleaved protein using standard chromatography techniques. The reference S protein sequence was from strain hCoV-19/Wuhan/IVDC-HB-01/2019 (EPI_ISL_402119). The isolated RBD, residues 328 to 541 of the S protein, was fused to a linker sequence containing a TEV-protease recognition site, followed by a human IgG1 Fc sequence. Fc-fusions of either the reference sequence, or containing mutations, were expressed in CHO cells and purified. The extracellular domain (ECD, residues 18 to 618) of ACE2 was expressed in CHO cells as an Fc-fusion protein containing a TEV-protease recognition site. Monomeric ACE2 ECD was generated by TEV protease digestion and purification of the ACE2 using standard chromatography techniques. For protein crystallization experiments, the isolated receptor binding domain (RBD, residues 329 to 527 of the S protein), was fused to a 6-His tag at the C-terminus, expressed in CHO cells, captured by Ni+-based immobilized metal affinity chromatography, enzymatically de-glycosylated using endoglycosidase-H, and purified by cation exchange chromatography. The Fab portion of LY-CoV1404, containing mutations in the constant region known to encourage crystallization, LY-CoV1404-CK Fab (Lieu et al., 2020a), was expressed in CHO cells, and purified by standard chromatography techniques. The Fab:RBD complex was prepared by mixing the components with 20% molar excess of the RBD, and then the complex purified from the excess RBD by size-exclusion chromatography.

### Method details

#### Single-cell screening and recovery

A blood sample from a 35-year-old individual hospitalized with severe COVID-19 disease was obtained in early-2020, approximately 60 days following the onset of symptoms. PBMC samples were collected under institutional review board (IRB)-approved protocols as part of the Hospitalized and Ambulatory Adults with Respiratory Viral Infections (HAARVI) study at the University of Washington (protocol #STUDY00000959) and Vaccine Research Center (VRC), National Institute of Allergy and Infectious Diseases (NIAID) and National Institutes of Health (NIH; protocol-VRC400, NIH-07IN194). Cells were thawed, activated in culture to generate antibody secreting cells and enriched for B cell lineage cells prior to injection into AbCellera’s microfluidic screening devices with either 91,000 or 153,000 individual nanoliter-volume reaction chambers. Single cells secreting target-specific antibodies were identified and isolated using three assay types: a multiplexed bead assay using multiple optically-encoded beads, each conjugated to the soluble pre-fusion stabilized S protein of either SARS-CoV-2 or WIV1 S with T4-foldon domain, 3C protease cleavage site, 6x His-tags, and twin-strep tags, the SARS-CoV-2 S1 subunit or negative controls (bovine serum albumin [BSA] His-tag and T4-foldon trimerization domain), and a live cell assay using passively dyed suspension-adapted Chinese hamster ovary (CHO) cells transiently transfected to surface-express full-length SARS-CoV-2 S protein (GenBank ID MN908947.3) with a green fluorescent protein (GFP) reporter, and non-transfected cells as a negative control. For the soluble assay, the IgG secreted by B-cells was captured on beads using the constant region. Binding to secreted IgG immobilized onto beads was subsequently assessed using soluble fluorescently labeled SARS-CoV-2 S or SARS-CoV-2 RBD antigen.

Beads or cells were flowed onto microfluidic screening devices and incubated with single antibody-secreting cells, and mAb binding to cognate antigens was detected via a fluorescently labeled anti-human IgG secondary antibody, or soluble antigen labeled with fluorophore. Positive hits were identified using machine vision and recovered using automated robotics-based protocols.

#### Sequencing, bioinformatic analysis, and cloning

Single-cell polymerase chain reaction (PCR) and custom molecular biology protocols generated NGS sequencing libraries (MiSeq, Illumina) using automated workstations (Bravo, Agilent). Sequencing data were analyzed using a custom bioinformatics pipeline to yield paired heavy and light chain sequences for each recovered antibody-secreting cell (Jones et al., 2020). Each sequence was annotated with the closest germline (V(D)J) genes, degree of somatic hypermutation, and potential sequence liabilities. Antibodies were considered members of the same clonal family if they shared the same inferred heavy and light V and J genes and had the same CDR3 length. The variable (V(D)J) region of each antibody chain was PCR amplified and inserted into expression plasmids using a custom, automated high-throughput cloning pipeline. Plasmids were verified by Sanger sequencing to confirm the original sequence previously identified by NGS. Antibodies were recombinantly produced by transient transfection in either human-embryonic kidney (HEK293) or CHO cells.

#### Epitope binning

All epitope binning and ACE-2 blocking experiments were performed on a Carterra® LSA™ instrument equipped with an HC-30M chip type (Carterra-bio), using a 384-ligand array format as previously described (Shu and McCauley, 2017). For epitope binning experiments, antibodies coupled to the chip surface were exposed to various antibody:antigen complexes. Samples were prepared by mixing each antibody in 10-fold molar excess with antigen (1:1 freshly prepared mix of 400 nM antibody and 40 nM antigen, both diluted in 1X HBSTE + 0.05% BSA running buffer). Each antigen-antibody premix was injected sequentially over the chip surface for 5 minutes (association phase to ligand printed onto chip previously), followed by a running buffer injection for 15 minutes (dissociation phase). Two regeneration cycles of 30 seconds were performed between each premix sample by injecting 10 mM glycine pH 2.0 onto the chip surface. An antigen-only injection (20 nM concentration in the running buffer) was performed every 8 cycles to assess maximum binding to S protein and in-order to accurately determine the binning relationship.

The data were analyzed using the Carterra Epitope analysis software for heat map and competition network generation. Analyte binding signals were normalized to the antigen-only binding signal, such that the antigen-only signal average is equivalent to one RU (response unit). A threshold window ranging from 0.5 RU to 0.9 RU was used to classify analytes into 3 categories: blockers (binding signal under the lower limit threshold), sandwichers (binding signal over the higher limit threshold) and ambiguous (binding signal between limit thresholds). Antibodies with low coupling to the chip, poor regeneration or with absence of self-blocking were excluded from the binning analysis. Like-behaved antibodies were automatically clustered to form a heat map and competition plot.

To test the antibodies’ ability to block ACE2, antibodies coupled to the HC-30M chip as described above were exposed to SARS-CoV-2 S protein:ACE2 complex. A freshly prepared 1:1 mix of 40 nM of SARS-CoV-2 S protein and 400 nM of untagged ACE2, both diluted in HBSTE + 0.05% BSA, were tested for binding to the immobilized mAbs on the prepared HC-30M chips, with association for 5 minutes and dissociation for 5 minutes. A SARS-CoV-2 S protein injection at 20 nM was included to assess for maximum binding, as well as a ACE2 injection at 200 nM to assess for non-specific binding. Regeneration was performed in 20 mM glycine pH 2.0 with 1M NaCl for 30 seconds twice.

#### Single-cycle kinetics

Surface plasmon resonance (SPR) capture experiments were performed on a Biacore 8K instrument equipped with a SA chip type (Cytiva, USA). The instrument uses one microfluidic module, an 8 multi-flow channel, to deliver samples to the chip surface via a unidirectional flow of sample at a set flow rate and concentration. The chip contains 8 flow cells, i.e. up to 8 ligands can be captured and analyzed at the same time. Streptavidin is pre-coated on the chip by the manufacturer.

Single cycle kinetics of LY-CoV1404 in Fab format was performed using the SA chip on the Biacore instrument as described herein. The chip was pre-conditioned with 3 injections of 10 mM NaOH buffer for 30 sec each, at a flow rate of 10 uL/min. The antigen of interest displaying a Strep tag was diluted to 5 nM in HBS-EP+ buffer (10 mM HEPES, 150 mM NaCl, 3 mM EDTA and 0.05% v/v Surfactant P20), then flowed over the SA chip for 60 sec at a flow rate of 10 uL/min. Fab LY-CoV1404 was diluted in HBS-EP+ buffer to various concentrations (300 nM, 100 nM, 33.3 nM, 11.1 nM, 3.7 nM, 1.2 nM). Each concentration was then serially flown over, from lowest concentration to highest, starting with a buffer (blank) injection, for 120 sec at a flow rate of 30 uL/min. After the Fab injections, HBS-EP+ buffer was injected for 600 sec at a flow rate of 30 uL/min. Finally, the chip was regenerated with a single injection of 10 mM Glycine, pH 2.0 for 30 s at a flow rate of 30 uL/min.

The data were analyzed using the Biacore Insight Evaluation Software and the pre-defined single cycle kinetics method. The curves were reference and blank subtracted and then fit to a 1:1 Langmuir binding model to generate association (ka) and dissociation (kd) kinetic rate constants and binding affinity constants (KD).

#### SPR affinity measurements

A Carterra® LSA™ instrument was used to measure binding kinetics of LY-CoV1404 to SARS-CoV-2 S protein, D614G S protein variant, RBD (receptor-binding domain) protein and mutant RBDs. Assays were performed according to the manufacturer’s operational guidelines. The instrument used a multi-channel buffer of 25 mM 2-(N-morpholino) ethanesulfonic acid (MES), pH 5.5, and a single-channel buffer of 10 mM 2-[4-(2-hydroxyethyl)piperazin-1-yl]ethanesulfonic acid (HEPES), 150 nM NaCl, 3 mM EDTA, and 0.05% v/v surfactant P20 (HBS-EP+).

To investigate binding of LY-CoV1404 mAb to SARS-CoV-2 S protein, D614G S protein variant and RBD protein, a HC30M chip was used, and the array preparation was performed as described above, coupling antibody diluted to 3 µg/mL in 10 mM acetate, pH 4.0, for 10 minutes, and deactivation for 7 minutes in 1 M ethanolamine, pH 8.5.

To measure binding kinetics and affinity, the mAb-coupled HC30M chip surface was exposed to injections of the proteins, with an association period of 5 minutes and dissociation period of 15 minutes. The tested concentrations of the trimeric S proteins were 300, 100, 33.3, 11.1, 3.70, 1.23, and 0.41 nM in HBS-EP+ containing 0.1 mg/mL BSA. The tested concentrations of the RBD protein were 400, 100, 25, 6.25 and 1.56 nM in HBS-EP+ containing 0.1mg/mL BSA. Regeneration of the chip surface between the different concentrations was performed using 20 mM glycine, pH 2.0, for 30 seconds twice. Kinetic data was analyzed using Carterra KIT™ software using a 1:1 Langmuir binding model.

To investigate binding of LY-CoV1404 Fab to reference and mutant RBDs a HC30M chip was used, and the array preparation was performed as described above, coupling Fc-fused RBD proteins diluted 5 µg/mL and 10 µg/mL in 10 mM acetate, pH 4.0, for 10 minutes, and deactivation for 7 minutes in 1 M ethanolamine, pH 8.5.

To measure binding kinetics and affinity, the RBD-coupled HC30M chip surface was exposed to injections of LY-CoV1404 Fab, with an association period of 5 minutes and dissociation period of 15 minutes. The tested concentrations of LY-CoV1404 Fab were 300, 100, 33.3, 11.1, 3.70, 1.23, and 0.41 nM in HBS-EP+ containing 0.1 mg/mL BSA. Regeneration of the chip surface between the different concentrations was performed using 20 mM glycine, pH 2.0, for 30 seconds twice. Kinetic data was analyzed using Carterra KIT™ software using a 1:1 Langmuir binding model or steady state model for samples that resulted in a poor 1:1 binding fit. Mutations where binding was observed but neither model resulted in good fits were reported as having affinities greater than the highest concentration of analyte tested. The reported affinity values are averages of multiple measurements for most variants, but the number of replicates varied depending on the variant and ranged from 2 to 26 replicate measurements.

#### Multi-cycle kinetics

The capture molecule, an anti-human IgG (Fc) antibody, was immobilized on a Biacore CM5 chip by direct coupling. The chip surface was first activated by flowing a freshly prepared 1:1 activation mixture of 100 mM S-NHS, 400 mM EDC for 10 min at a flow rate of 10 uL/min. Anti-human IgG (Fc) antibody was diluted to 25 ug/mL in 10 mM Sodium Acetate buffer pH 4.5, then injected using all 8 channels, at a flow rate of 10 uL/min for 10 min. The chip was washed with HBS-EP+ (10 mM HEPES, 150 mM NaCl, 3 mM EDTA and 0.05% v/v Surfactant P20) for 1 min, at a flow rate of 30 uL/min. Finally, excess reactive esters were quenched by flowing 1 M ethanolamine for 10 min at a flow rate of 10 uL/min, followed by 3 conditioning steps of 30 s each, in 10 mM NaOH buffer, at a flow rate of 10 uL/min.

Multi cycle kinetics of LY-CoV1404 was performed using the previously prepared CM5 chip on the Biacore instrument as described herein. LY-CoV1404 was diluted to 5 nM in HBS-EP+ buffer (as above), then flowed over the CM5 chip for 30 sec at a flow rate of 10 uL/min. Each antigen of interest was diluted in HBS-EP+ buffer to various concentrations (100 nM, 33.3 nM, 11.1 nM, 3.7 nM, 1.2 nM). Each concentration was then serially flown over, from lowest concentration to highest, starting with a buffer (blank) injection, for 120 sec at a flow rate of 30 uL/min. After each antigen injection, HBS-EP+ buffer was injected for 600 sec at a flow rate of 30 uL/min. Then the chip was regenerated with a single injection of 3M MgCl2 for 30 s at a flow rate of 30 uL/min before the next antigen injection.

The data were analyzed using the Biacore Insight Evaluation Software. The curves were referenced and blank subtracted and then fit to a 1:1 Langmuir binding model to generate apparent association (ka) and dissociation (kd) kinetic rate constants and binding affinity constants (KD).

#### Cell-based binding assay

Unique antibody sequences were confirmed to bind the screening target (SARS-CoV-2 full length S protein) using high throughput flow cytometry. CHO cells were transiently transfected to express the full-length S protein of either wild type SARS-CoV or SARS-CoV-2 or mutant SARS-CoV-2 (R21I, T22I, T29I, H49Y, D138H, Q490E, N439K, G476S, S477N, T478I, V483A, F490S, S494P, N501Y, G504D, A520S, D614G, B.1.1.7, B.1.351) on the cell surface. Suspension CHO cells were transiently transfected with the plasmid using electroporation. Full length native conformation S protein expression was confirmed by testing with benchmark antibodies discovered against SARS-CoV that target different stalk and head domains using flow cytometry.

Purified antibodies at 50nM antibody concentration were incubated with the readout cells or an untransfected control CHO cells for 30 minutes at 4°C. CHO cells were washed, and binding was detected by using a fluorescently labeled anti-human secondary antibody. Fluorescence was measured using high throughput plate-based flow cytometry. Benchmark antibodies identified to bind to SARS-CoV were used as positive controls due to similarity in S protein sequences between SARS-CoV and SARS-CoV-2; human IgG isotype and an irrelevant antibody were used as negative controls. Median fluorescence intensity of each antibody was normalized over the median fluorescence intensity of the human isotype control for respective antigens. The median fold over isotype values from different validation experiments were plotted. Antibody values greater than 5-fold over isotype were considered as binders. The cut-off value was determined based on the binding to the negative controls.

#### Pseudovirus Production and Characterization

Mutagenesis reactions were performed using the QuickChange Lightning Site-Directed Mutagenesis Kit (Agilent Cat # 210519) using as template a S protein mammalian expression vector based on the Wuhan sequence (Genbank MN908947.3) with a deletion of the C-terminal 19 amino acids. Pseudoviruses bearing mutant S proteins were produced using the ΔG luciferase recombinant vesicular stomatitis virus (rVSV) system (KeraFast EH1025-PM)(Whitt, 2010). Briefly, 293T cells were transfected with individual mutant spike expression plasmids, and 16 to 20 hours later, transfected cells were infected with VSV-G-pseudotyped ΔGluciferase rVSV. Sixteen to 20 hours following infection, conditioned culture medium was harvested, clarified by centrifugation at 1320 x g for 10 minutes at 4°C, aliquoted, and stored frozen at -80°C. Infectious titers were determined for a subset of virus preparations by infection of Vero-E6 cells (ATCC CRL-1586) with serially diluted virus followed by staining with an anti-luciferase antibody (Novus Cat # NB600-307PEATT594) and analysis by fluorescence-activated cell sorting using a Becton Dickinson LSRFortessaTM X-20. Particle titers were determined by isolation of genomic RNA and quantitation by RT-qPCR (NEB # E3031) using an Applied Biosystems vii7A™ Real-Time PCR system. Relative luciferase reporter signal read-out was determined by luciferase assay (Promega Cat # E2650) of extracts from VeroE6 cells infected with serially diluted virus. Luciferase activity was measured on a PerkinElmer EnVision 2104 multilabel reader.

#### Pseudovirus Neutralization Assays

Neutralization assays were carried out as described (Nie et al., 2020). Virus preparation volumes were normalized to equivalent signal output (RLU, relative light units) as determined by luciferase activity following infection with serially diluted virus. Eleven-point, 2-fold titrations of LY-CoV1404 were performed in 96-well plates in duplicate and pre-incubated with a fixed amount of pseudovirus for 20 minutes at 37°C. Following pre-incubation, the virus-antibody complexes were added to 20,000 VeroE6 cells/well in white, opaque, tissue culture-treated 96 well plates, and incubated 16 to 20 hours at 37°C. Control wells included virus only (no antibody, quadruplicate) and cells only (duplicate). Following infection, cells were lysed, and luciferase activity was measured.

#### Pseudotyped lentivirus Neutralization Assays

SARS-CoV-2 spike pseudotyped lentiviruses that harbor a luciferase reporter gene were produced and neutralization assay was performed as described previously (Corbett et al., 2020; Naldini et al., 1996). Pseudovirus was produced by co-transfection of 293T cells with plasmids encoding the lentiviral packaging and luciferase reporter, a human transmembrane protease serine 2 (TMPRSS2), and SARS-CoV-2 S (Wuhan-1, Genbank #: MN908947.3; D614G and B.1.1.529 containing A67V, H69-, V70-, T95I, G142D, V143-, Y144-, Y145-, N211-,L212I, ins214EPE, G339D, S371L, S373P, S375F, K417N, N440K, G446S, S477N, T478K, E484A, Q493R, G496S, Q498R, N501Y, Y505H, T547K, D614G, H655Y, N679K, P681H, N764K, D796Y, N856K, Q954H, N969K, L981F amino acid changed compared to WA-1) or S variant genes (Wang et al., 2021a) using Lipofectamine 3000 transfection reagent (ThermoFisher, CA). Forty-eight hours after transfection, supernatants were harvested, filtered and frozen. For neutralization assay serial dilutions (2 dilutions at 10 and 1 µg/ml for the initial screen assay or 8 dilutions for the full curve at 10-0.0006 µg/ml) of monoclonal antibodies were mixed with titrated pseudovirus, incubated for 45 minutes at 37 °C and added to pre-seeded 293T-ACE2 cells (provided by Dr. Michael Farzan) or 293 flpin-TMPRSS2-ACE2 cells (made by Adrian Creanga, VRC, NIH) in triplicate in 96-well white/black Isoplates (Perkin Elmer). Following 2 hours of incubation, wells were replenished with 150 µL of fresh medium. Cells were lysed 72 hours later, and luciferase activity (relative light units, RLU) was measured. Percent neutralization was normalized considering uninfected cells as 100% neutralization and cells infected with only pseudovirus as 0% neutralization. Neutralization IC50, IC80 and IC90 titers were calculated using GraphPad Prism 8.0.2.

#### Identification of escape mutation positions

The potential for SARS-CoV-2 to develop resistance under selective pressure of LY-CoV1404, was studied using in vitro directed evolution. The RBD comprised of residues 319 to 541 of SARS-CoV-2 S protein (Wuhan sequence, Genbank MN908947.3) was expressed on the surface of yeast cells. Surface-bound RBD was confirmed to be reactive with recombinant soluble hACE2 and LY-CoV1404 by flow cytometry. Soluble hACE2 used for *in vitro* selection screens (Agro BioSciences Cat # AC2-H82E6) comprised residues 18 to 634 of Uniprot Q9BYF1 and contained a C terminus Avi tag to enable fluorescence after binding to streptavidin labeled with phycoerythrin. A library was constructed in which all amino acid residues were sampled at positions 331 to 362 and 403 to 515 of the spike glycoprotein. RBD variants that could still bind 4 nM hACE2 in the presence of antibody at 5 nM (14-fold IC50) concentration were isolated after multiple rounds of selection by fluorescence activated cell sorting and sequenced. Detailed characterization of isolated variants was limited to clones that contained only single amino acid changes and that could occur by a single nucleotide change within the codon to better reflect errors that can arise during viral replication in vivo. Additionally, clones containing specific variants of interest observed from public databases and/or shown to be resistant with other authorized monoclonal antibodies were analyzed.

LY-CoV1404 inhibition of binding to soluble hACE2 used a flow cytometry assay to calculate IC50 for RBD variants of interest expressed on the surface of yeast. The in vitro inhibition assay used 1 nM soluble hACE2 incubated with increasing concentrations of LY-CoV1404, ranging from 0 to 3.75 ug/mL with each yeast RBD variant of interest. Soluble hACE2 binding was performed at room temperature for 20 minutes followed by transfer of reactions to ice for another 10 minutes to quench dissociation. Fluorescence was measured after addition of streptavidin-phycoerythrin for 20 minutes on ice followed by multiple washes with cold buffer. Mean fluorescence intensity (MFI) was normalized for each concentration response curve (CRC) using the maximum MFI at no mAb inhibitor (MAX) and the background MFI (Bgnd). Percent inhibition of S protein binding to ACE2 was defined as follows:

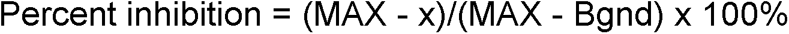

where x was the MFI at the tested concentration of mAb. CRCs were fit with a four-parameter logistic function. All four parameters were estimated from the fitting. Absolute IC50 (50% absolute inhibition) values were reported. Standard error and 95% confidence intervals for IC50 estimates are reported and, in some cases, a fixed top (Top = 100) was used to stabilize the standard error estimate. IC50 ratios between mutant S protein and wild type (WT) were reported. For mutant S proteins that exhibited no mAb inhibition, fold-change was reported at 1x of the highest concentration of mAb tested. For any single experimental batch, inhibition data were obtained in duplicate or triplicate. WT RBD was analyzed with every experiment and the consensus estimate as of the date indicated is reported. If select mutants of interest were analyzed in more than one experimental batch, then the geometric mean is reported.

#### Authentic SARS-CoV-2 Neutralization

Work with authentic SARS-CoV-2 at USAMRIID was completed in BSL-3 laboratories in accordance with federal and institutional biosafety standards and regulations. Vero-76 cells were inoculated with SARS-CoV-2 (GenBank MT020880.1) at a MOI = 0.01 and incubated at 37°C with 5% CO2 and 80% humidity. At 50 h post-infection, cells were frozen at -80°C for 1 h, allowed to thaw at room temperature, and supernatants were collected and clarified by centrifugation at ∼2,500 x g for 10 min. Clarified supernatant was aliquoted and stored at -80°C. Sequencing data from this virus stock indicated a single mutation in the S glycoprotein (H655Y) relative to Washington state isolate MT020880.1.

Authentic SARS-CoV-2 plaque reduction assays were also conducted in BSL3 laboratories at UTMB. Three natural isolates were used to measure mAb neutralization: USA/WA/1/2020 (BEI resources number NR52281), Italy-INMI1 (European Virus Archive – Global, Ref #008V03893), and the Netherlands (1363454/NL/2020). The NL isolate is known to carry the D614G S protein variation. For studies conducted with variant viruses, virus isolates were constructed using reverse genetics (in the case of the rWA1, rWA1 E484K, and rWA1 E484Q viral isolates), or from patient isolates expanded in cell culture (in the case of the WA1, B.1.1.7, and B.1.351 viral isolates). Methods for the recombinant virus generation are described in detail in a recent publication (Xie et al. 2021). In brief, a viral cDNA encoding the WA strain of SARS-CoV-2 was designed to express authentic virus containing spike genes corresponding to either wild-type, E484K or the E484Q sequence. For naturally occurring variants, four clinical isolates were used as the inocula: USA/WA/1/2020 (BEI Resources NR52281), B.1.351 (USA/MD-HP01542/2021, UTMB WRCEVA collection), B.1.1.7 (USA/CA_CDC_5574, UTMB WRCEVA collection) and B.1.617.2 (GNL-751-TVP23523, UTMB WRCEVA collection). All three variant strains were characterized by NGS sequencing at UTMB to confirm the expected spike protein mutations: B.1.1.7 contained Δ69-70/Δ145/N501Y/A570D/D614G/P681H/R682Q/T716I/S982A/D1118H; B.1.351 contained L18F/D80A/D215G/Δ242-244/K417N/E484K/N501Y/D614G/A710V; and B.1.617.2 contained T19R/G142D/Δ156-157/R158G/A222V/L452R/T478K/D614G/P681R/R685H/D950N/V1264L. Recombinant virus isolates (in the case of the rWA1, rWA1 E484K, and rWA1 E484Q viral isolates) were constructed using reverse genetics as previously described (Xie et al., 2021b). Virus stocks were grown by inoculating cultured Vero E6 cells, followed by incubation at 37°C until cytopathic effects (CPE) were evident (typically 48 to 72 hours). Expansion was limited to 1 to 2 passages in cell culture to retain integrity of the original viral sequence. The virus stock was quantified by standard plaque assay, and aliquots were stored at -80°C. A freshly thawed aliquot was used for each neutralization experiment.

#### Authentic SARS-CoV-2 IFA neutralization assay

A pre-titrated amount of authentic SARS-CoV-2/MT020880.1, at final multiplicity of infection of 0.2, was incubated with serial dilutions of monoclonal antibodies for 1 h at 37°C. The antibody-virus mixture was applied to monolayers of Vero-E6 cells in a 96-well plate and incubated for 1 hour at 37°C in a humidified incubator. Infection media was then removed, and cells were washed once with 1X PBS, followed by addition of fresh cell culture media. Culture media was removed 24 hours post infection and cells were washed once with 1X PBS. PBS was removed and plates were submerged in formalin fixing solution, then permeabilized with 0.2% Triton-X for 10 minutes at room temperature and treated with blocking solution. Infected cells were detected using a primary detection antibody recognizing SARS-CoV-2 nucleocapsid protein (Sino Biological) following staining with secondary detection antibody (goat α-rabbit) conjugated to AlexaFluor 488. Infected cells were enumerated using the Operetta high content imaging instrument and data analysis was performed using the Harmony software (Perkin Elmer).

#### PRNT assay

Vero-E6 cells were seeded in a 24-well plate 48 hours before the assay (Figure 2). Seventy-five plaque forming units (pfu) of infectious clone hCoV-19/Canada/ON_ON-VIDO-01-2/2020 were mixed with serial dilutions of monoclonal antibodies and incubated at 37°C for 60 minutes. Virus and antibody mix was added to each well and incubated for 1 h in a 37°C + 5% CO2 incubator with rocking every 10-15 min. Plaque assay media (complete MEM media with 1% BGS + 1% low melting point agarose) was overlaid on top of the inoculum and incubated at 37°C + 5% CO2 incubator for 48 hours. For plaque visualization, an MEM-Neutral Red overlay was added on day 2 and plaques counted manually on day 3 or day 4.

Plaque Reduction Neutralization test assays were performed in 6-well plates (Table 2). Vero E6 cells were seeded at a concentration of approximately 106 cells/well and grown overnight at 37°C in 5% CO2 to reach 95% confluency. The next day, serial three-fold dilutions of LY-CoV1404 or bamlanivimab were prepared in Eagle’s minimal essential medium, mixed with approximately 100 pfu of SARS-CoV-2, and incubated for 1 to 2 hours on ice or at 37°C. The mAb/virus mixture was inoculated directly onto the cells (in duplicate wells) and allowed to adsorb for 1 hour at 37°C with 5% CO2, with rocking at 15-minute intervals. An overlay media composed of 1.25% Avicel RC 581 (FMC BioPolymer) in Eagles minimum essentials medium (MEM) with 5% FBS was added, and plates were incubated for 48 hours at 37°C with 5% CO2 for virus plaques to develop. After incubation, overlays were removed by aspiration and the cells were fixed with 10% buffered formalin containing crystal violet stain for 1 hour. Plaques were counted manually, and plaque forming units were determined by averaging technical replicates per sample. Percent neutralization was determined relative to virus-only control-treated samples.

#### Protein Crystallography

A 10 mg/mL solution of LY-CoV1404 Fab (with CrystalKappa mutations (Lieu et al., 2020b)) in complex with RBD was set up in vapor diffusion sitting drops at a ratio of 1:1 with a well solution of 100 mM Tris HCl pH 6.5, 20 % PEG MME 2K and 200 mM Trimethylamine N-oxide. Crystals appeared within two days, grew to their full size and were harvested on the fifth day after the set up. Crystals were flash-frozen in liquid nitrogen following 1-minute incubation in cryoprotectant solution containing mother liquor supplemented with additional 5% PEG MME 2K.

Diffraction data were collected at Lilly Research Laboratories Collaborative Access Team (LRL-CAT) and beamline at Sector 31 of the Advanced Photon Source at Argonne National Laboratory, Chicago, Illinois. Crystals stored in liquid nitrogen were mounted on a goniometer equipped with an Oxford Cryosystems cryostream maintained at a temperature of 100 K. The wavelength used was 0.9793 Å collecting 900 diffraction images at a 0.2 degree oscillation angle and 0.1 seconds exposure time on a Pilatus3 S 6M detector at a distance of 385 mm. The diffraction data were indexed and integrated using autoPROC (Vonrhein et al., 2011)/XDS (Kabsch, 2010) and merged and scaled in AIMLESS (Evans and Murshudov, 2013) from the CCP4 suite (Winn et al., 2011). Non-isomorphous data readily yielded initial structures by molecular replacement using the Fab portion of crystal structures from the proprietary Eli Lilly structure database and the SARS-CoV-2 S protein RBD from the public domain structure with the access code 6yla (Huo et al., 2020). The initial structure coordinates were further refined using Buster (Smart et al., 2012) applying isotropic temperature factors. Model building was performed with Coot (CCP4) and final structure validation with MolProbity (Chen et al., 2010) and CCP4 validation tools.

Structure superposition with other published Fab:S complexes, contact surface analysis, and supplemental figure generation was performed with MOE (Molecular Operating Environment, 2019.0101; Chemical Computing Group ULC), and detailed contact analysis with CCP4 CONTACT(Winn et al., 2011) and custom shell/Perl scripts.

#### Protein expression

A stabilized version of the SARS-CoV-2 S protein was employed for all studies (Hsieh 2020). The sequence corresponding to the B.1.1.7 variant extracellular domain (ECD) was acquired from the GSAID database. The stabilized construct consists of the residues of the SARS-CoV2 S protein 1-1208 with proline substitutions at position 814, 889, 896, 939, 983, 984, GSAS substitution at the furin cleavage site (679-682), C - terminal foldon trimerization domain, and an octa histidine tag. The cDNA sequence synthesized and cloned into pcDNA3.4 for mammalian cell expression (GenScript).

Fabs were generated by subcloning the variable region and CH1 of the heavy chain of LY-CoV1404 into a pcDNA based vector with C terminal 6-Histidine tag. The boundary of the CH1 domain was set according to the consensus sequence defining the beginning of the hinge region directly following CH1.

#### Protein purification and characterization

The plasmid encoding the B.1.1.7 stabilized S protein ECD sequence was transiently transfected into expi293 cells according to the manufacturer’s specifications. Briefly, 1 µg/mL plasmid DNA diluted in optiMem was mixed with the recommended expifectamine/ OptiMem solution and incubated for 15 min. The DNA/expifectamine mixture was added to expi293 cells diluted to 3X10e6 cells/mL. After 24 hours, the recommended volumes of enhancer 1 and enhancer 2 were added to the cells, which were harvested 5 days post transfection. The supernatant was separated from the cells via centrifugation at 3000 X g and further clarified by passage through a 0.22 um filter. Clarified supernatants were loaded onto a 5 mL Ni sepharose excel column (Cytiva), washed for 20 column volumes using PBS + 20 mM imidazole and eluted with 10 column volumes of PBS + 500 mM imidazole. The protein was immediately further purified using size exclusion chromatography using a superose 6 16/600 Hiload column (Cytiva). The protein was concentrated to 0.75 mg/mL (Amicon Ultra-15 100 kDa MWCO) and frozen in 100 µg aliquots for structural Biology studies.

Fab generation was initiated with plasmids encoding the truncated heavy chain and light chain of LY-CoV1404 (1 ug/mL each) that were co-transformed into expi293 cells (expifectamine, Gibco) for protein expression (5 days). Cell supernatants were collected, clarified and incubated with Ni-NTA beads to capture the Fab, followed by washing in PBS buffer (pH 7.4) and elution in the same buffer supplemented with 250 mM Imidazole.

#### Cryo electron microscopy

Purified B.1.1.7 S protein (0.8mg/ml) was mixed with LY-CoV1404 (0.5 mg/ml) and incubated on ice for 30 min. Quantifoil R 2/2, 300 mesh copper grids (Electron Microscopy Sciences) were plasma cleaned in a PELCO easiGlow Glow Discharge Cleaning System (Ted Pella). A 3μl drop of sample was applied to freshly glow discharged grids followed by vitrification in liquid ethane using Vitrobot Mark IV with a blot time of 30 seconds. A total of 5,202 micrographs were recorded on a 3000kV Titan Krios microscope (Thermo Fisher Scientific) equipped with a Falcon IV detector at a 130,000x nominal magnification corresponding to a pixel size of 0.98Å. The RELION program was used for particle picking, 2D and 3D class averaging (Scheres 2016).

#### GISAID Analysis

To estimate the incidence of SARS-CoV-2 variants with potential resistance to LY-CoV1404, we analyzed data from GISAID EpiCoV database (Shu and McCauley, 2017), focusing on recent data, i.e. genomes collected in a 3 month window from Dec 16 2020 to March 16 2021, and submitted prior to March 22 2021. This gave a total of 281240 samples in the dataset, of which 76053 were from North America. We aligned the genomes to the SARS-CoV-2 reference genome (Genbank file MN908947.3) using BWA-MEM (Li and Durbin, 2010), and performed variant calling and annotation using SAMTools (Danecek et al., 2021). All further analysis (i.e. slicing by time or geographic region) was performed using custom-written Python scripts.

#### Statistical Analysis

All the statistical details of specific experiments, which included the statistical tests used, number of samples, mean values, standard errors of the mean (SD), and p-values derived from indicated tests, had been described in the figure legends and showed in figures. Statistical analyses were conducted utilizing GraphPad Prism 8.0 or Microsoft Excel. Triplicate, sextuplicate, and other replicative data were presented as mean ± SD. Value of p < 0.05 was considered to be statistically significant and represented as an asterisk (∗). Value of p < 0.01 was supposed to be more statistically significant and described as double asterisks (∗∗). Value of p < 0.001 was considered the most statistically significant and represented as triple asterisks (∗∗∗). Value of p < 0.0001 was supposed to be extremely statistically substantial and described as quadruple asterisks (∗∗∗∗). For comparison between two treatments, a Student’s t-test was used. For comparison between each group with the mean of every other group within a dataset containing more than two groups, one-way ANOVA with Tukey’s multiple comparisons test was used.

## SUPPLEMENTARY FIGURES

**Figure S1.**
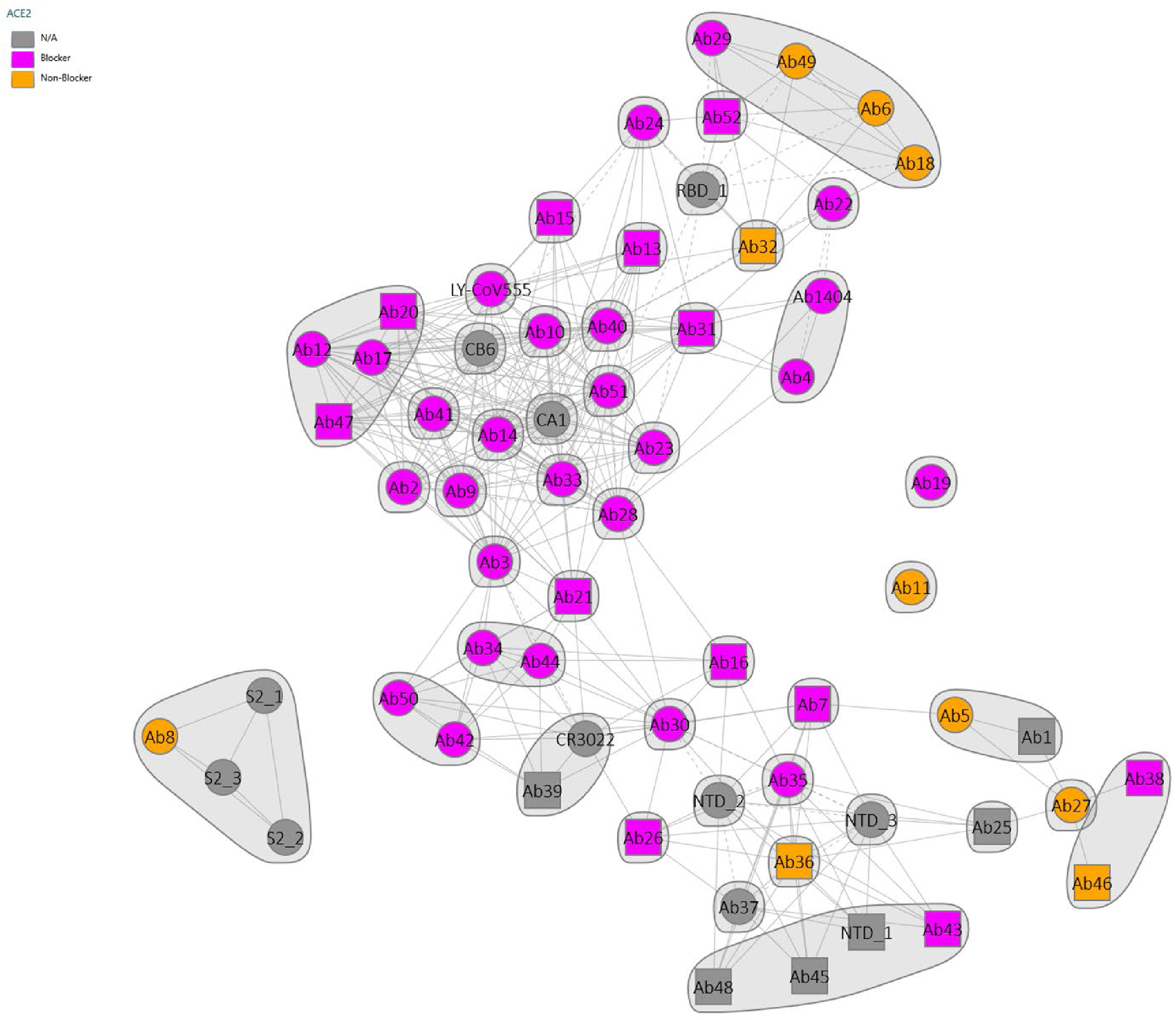
ACE2 competition with discovered antibodies. Competition plot of recombinantly expressed antibodies overlaid with ACE2 blocking information. ACE2 blockers and non-blockers are indicated by color (ACE2 blockers = Pink; ACE2 non-blockers = orange).

**Figure S2.**
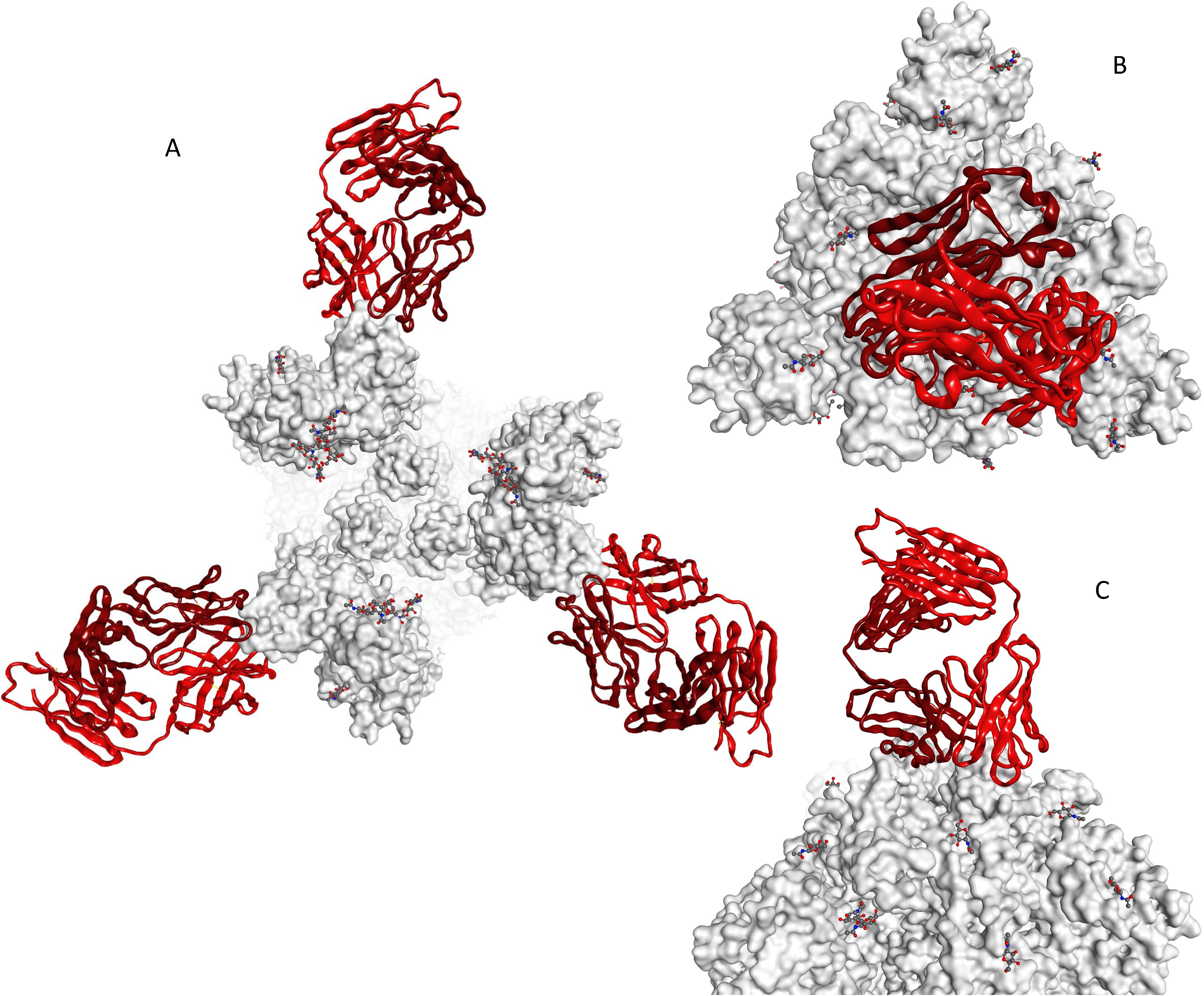
LY-CoV1404 binding to trimeric S protein. LY-CoV1404 modeled on 3 RBD-up (A) and 3 RBD-down (B, top; C, side) conformations of S protein. Only one molecule of LY-CoV1404 can bind to a 3-RBD-down spike trimer.

**Figure S3.**
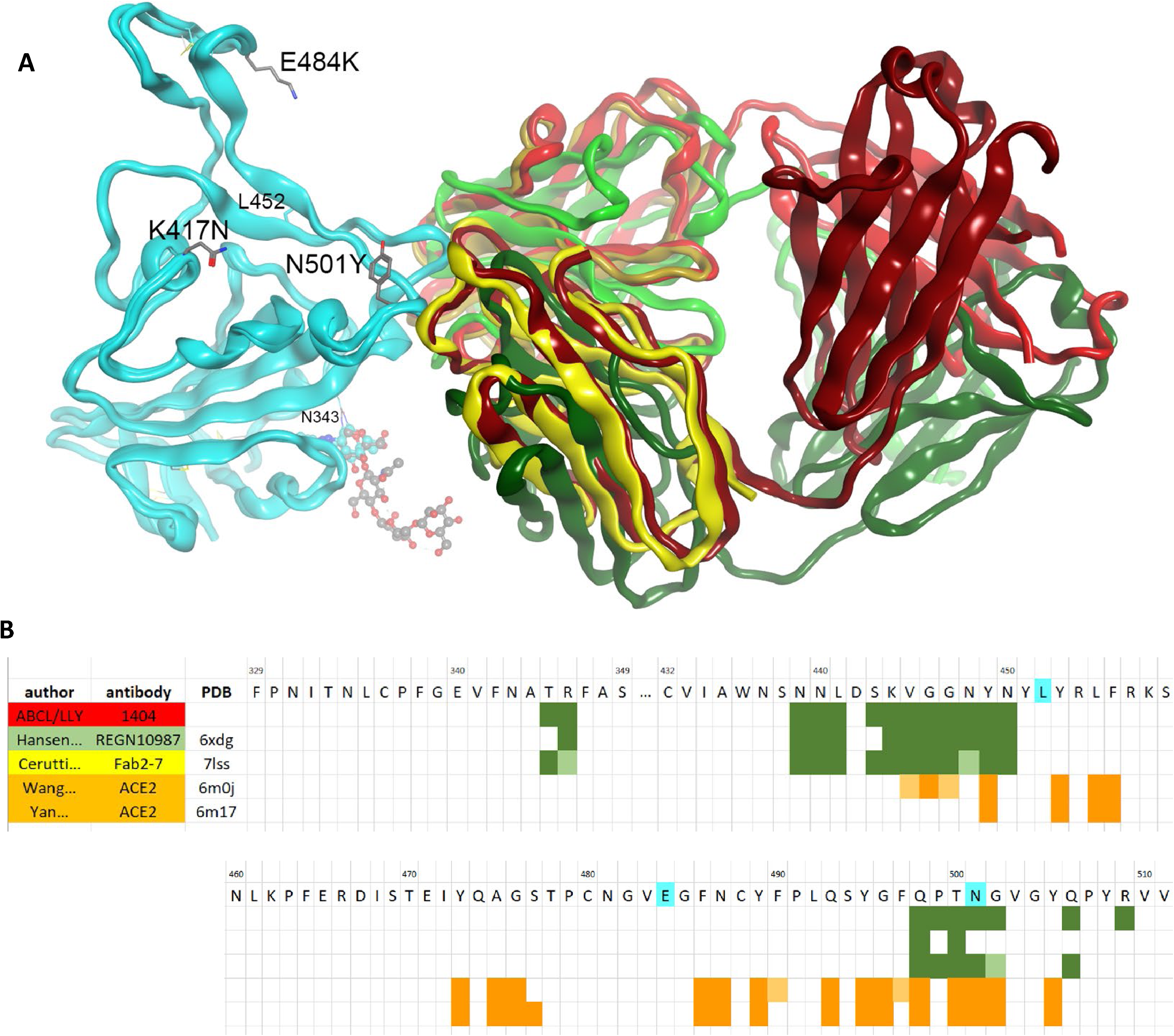
Antibody binding to variant B.1.351 and antibody epitopes. (A) Superposition of LY-CoV1404 (red), REGN10987 (6XDG, green), and Fab 2-7 (7LSS, yellow), with RBDs (cyan) from SARS-CoV-2 Wuhan-Hu-1 superposed on the B.1.351 variant (7NXA) (Dejnirattisai et al., 2021) showing the location of mutated sidechains. (B) Antibody and ACE2 epitopes highlighted on the RBD sequence.

**Table S1.**
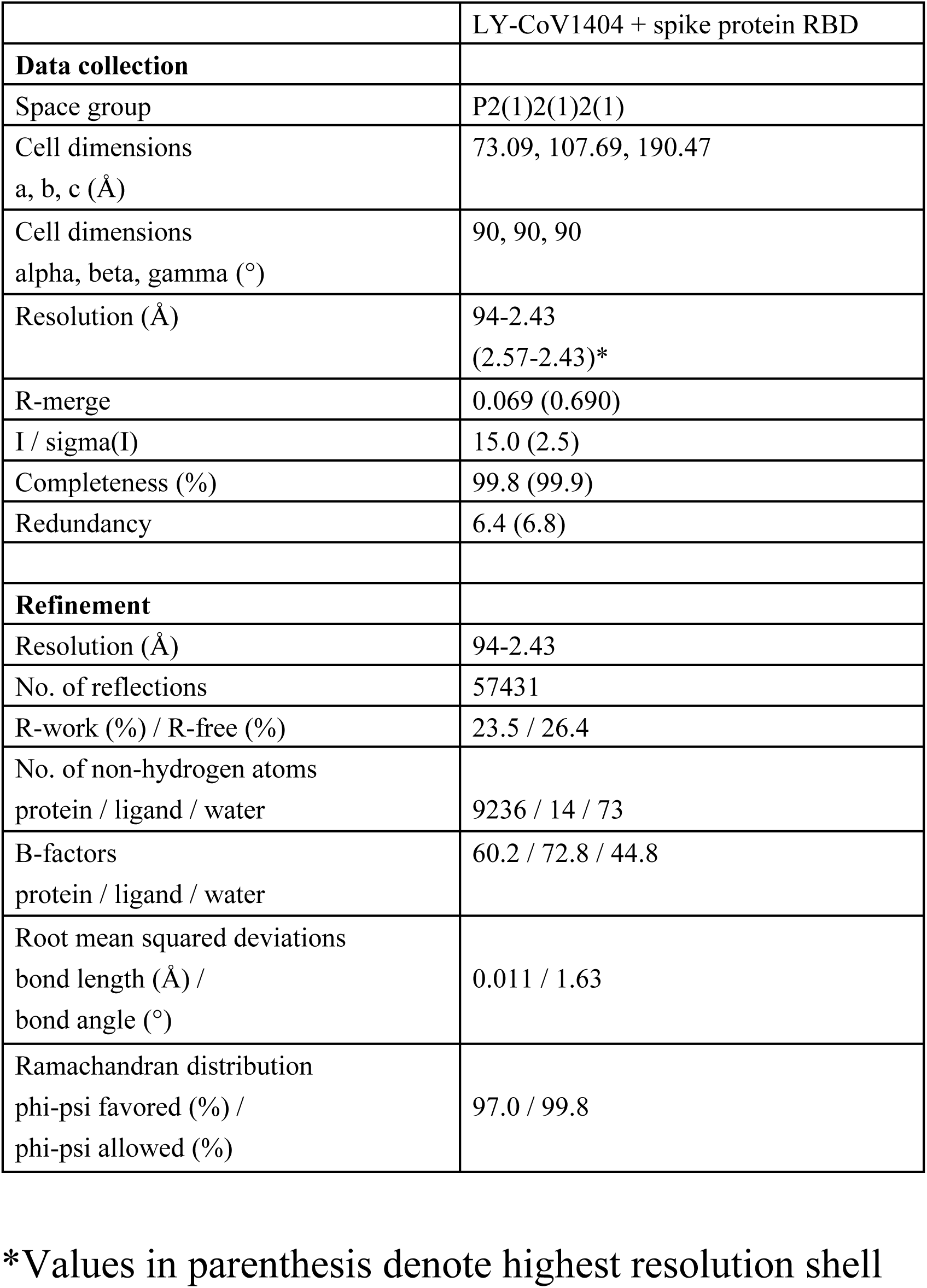
Crystallographic statistics

**Table S2.**
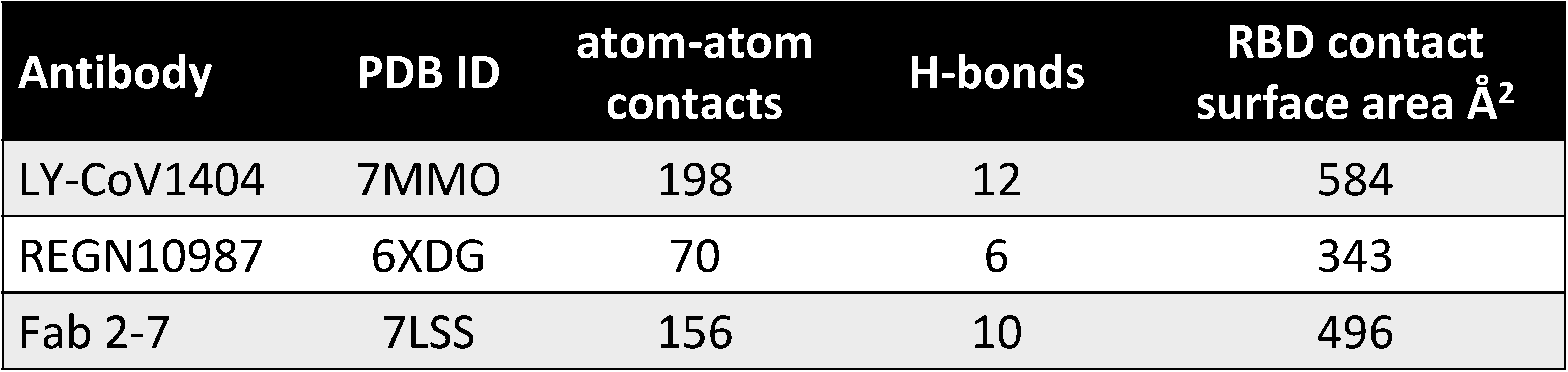
Summary of atomic interactions at the RBD epitope

