## Supplementary Materials for "LY-CoV1404 (bebtelovimab) potently neutralizes SARS-CoV-2 variants"

**Fig S1.** ACE2 Competition

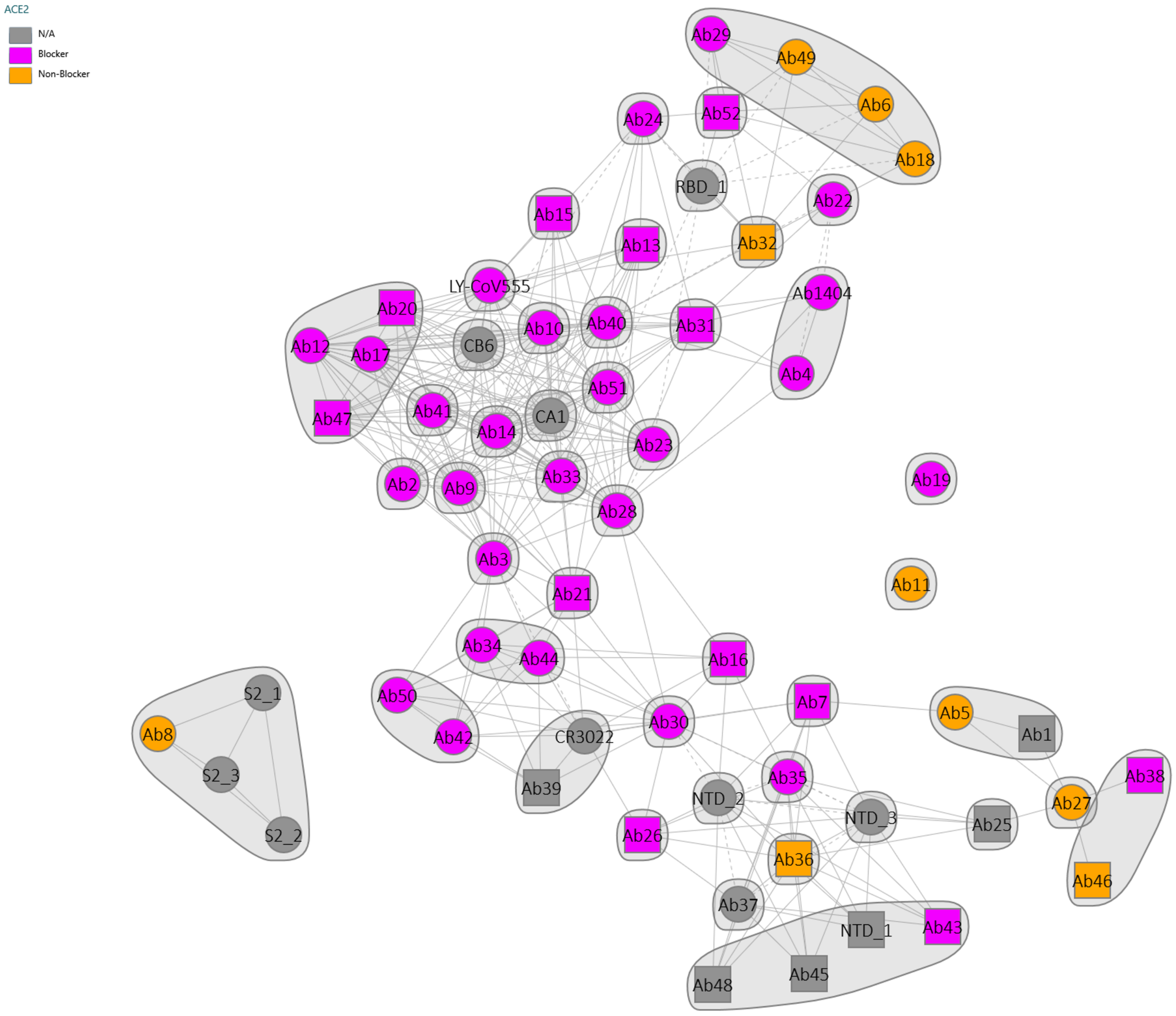

**Fig S2.** LY-CoV1404 binding to trimeric S protein

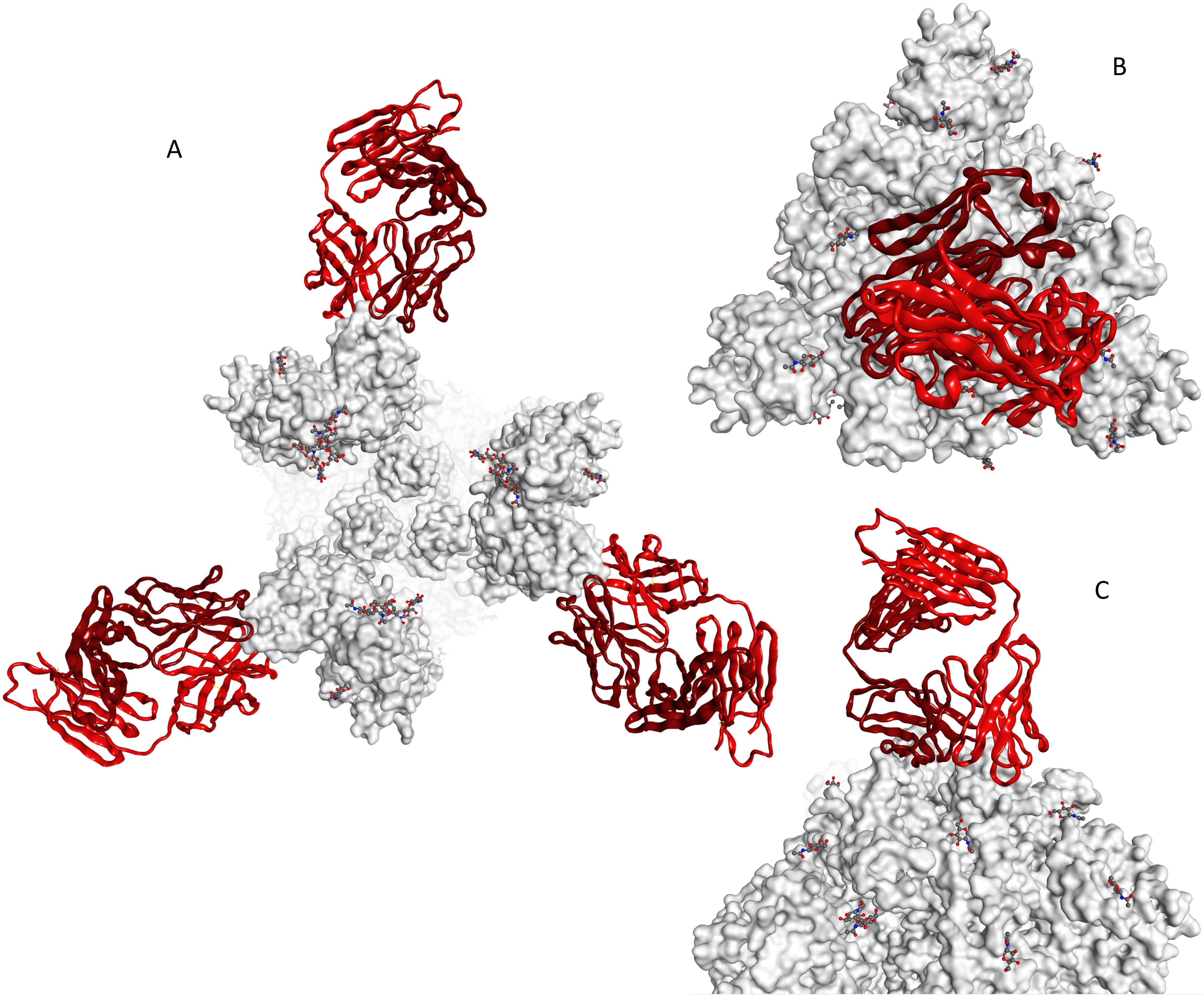

**Fig S3.** Antibody binding to variant B.1.351 and antibody epitopes.

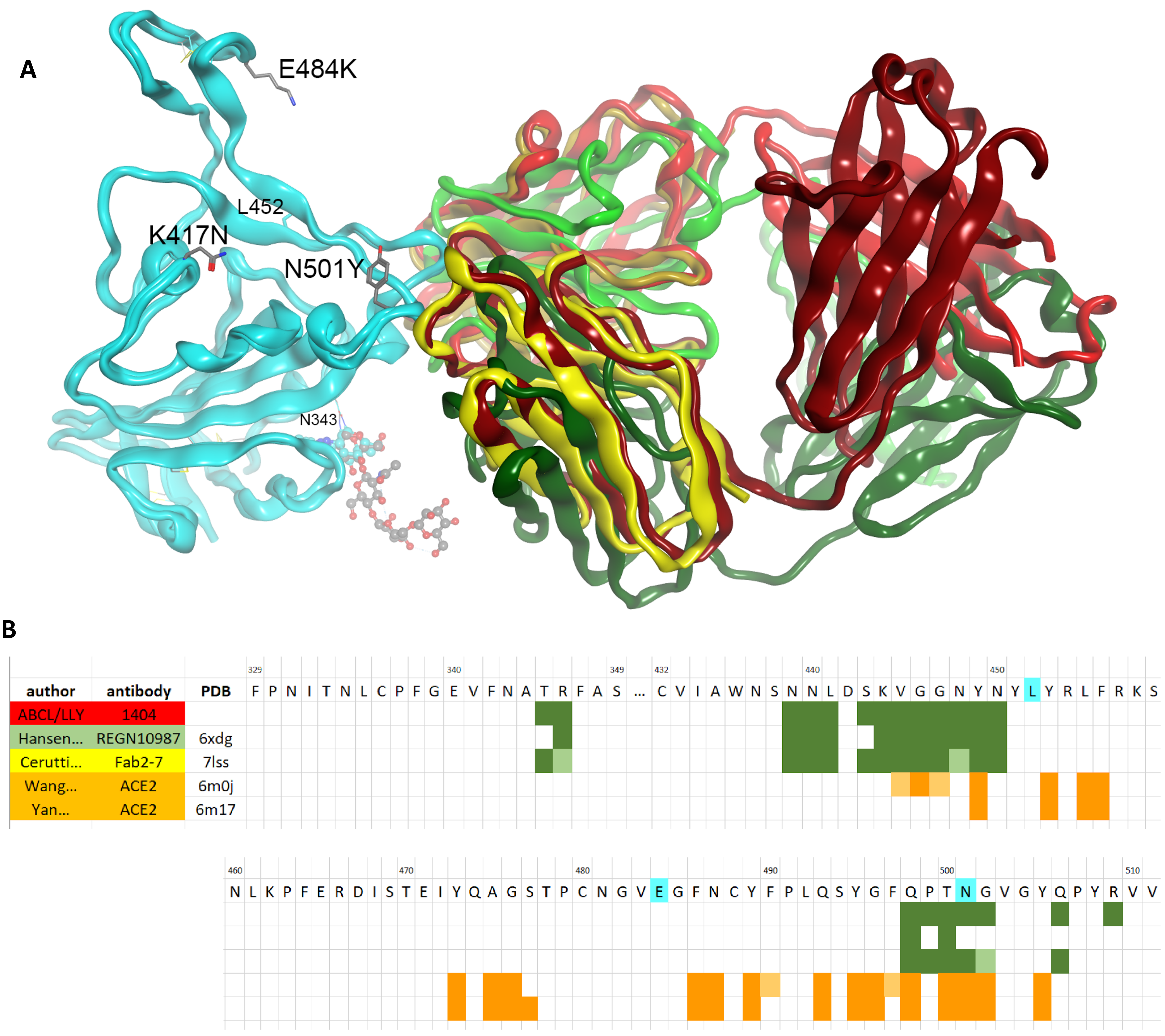

**Table S1.** Crystallographic statistics

| | LY-CoV1404 + spike protein RBD |
|---|---|
| **Data collection** | |
| Space group | P2(1)2(1)2(1) |
| Cell dimensions<br>a, b, c (Å) | 73.09, 107.69, 190.47 |
| Cell dimensions<br>alpha, beta, gamma (°) | 90, 90, 90 |
| Resolution (Å) | 94-2.43<br>(2.57-2.43)* |
| R-merge | 0.069 (0.690) |
| I / sigma(I) | 15.0 (2.5) |
| Completeness (%) | 99.8 (99.9) |
| Redundancy | 6.4 (6.8) |
| **Refinement** | |
| Resolution (Å) | 94-2.43 |
| No. of reflections | 57431 |
| R-work (%) / R-free (%) | 23.5 / 26.4 |
| No. of non-hydrogen atoms<br>protein / ligand / water | 9236 / 14 / 73 |
| B-factors<br>protein / ligand / water | 60.2 / 72.8 / 44.8 |
| Root mean squared deviations<br>bond length (Å) /<br>bond angle (°) | 0.011 / 1.63 |
| Ramachandran distribution<br>phi-psi favored (%) /<br>phi-psi allowed (%) | 97.0 / 99.8 |

*Values in parenthesis denote highest resolution shell

**Table S2.** Summary of atomic interactions at the RBD epitope.

| Antibody | PDB ID | atom-atom contacts | H-bonds | RBD contact surface area Å$^2$ |
|---|---|---|---|---|
| LY-CoV1404 | | 198 | 12 | 584 |
| REGN10987 | 6XDG | 70 | 6 | 343 |
| Fab 2-7 | 7LSS | 156 | 10 | 496 |
